# Double strand breaks drive toxicity in Huntington’s disease mice with or without somatic expansion

**DOI:** 10.1101/2025.05.27.654663

**Authors:** Aris A Polyzos, Ana Cheong, Jung Hyun Yoo, Lana Blagec, Zachary D Nagel, Cynthia T McMurray

## Abstract

There has been a substantial investment in elucidating the mechanism of expansion in hopes of identifying therapeutic targets for Huntington disease (HD). Although an expanded CAG allele is the causal mutation for HD, there is evidence that somatic expansion may not be the only disease driver. We report here that double strand breaks (DSBs) drive HD toxicity by an independent mechanism from somatic expansion. The mutant HD protein inhibits non-homologous end joining (NHEJ) activity, leading to the accumulation of DSBs. DSBs promote transcriptional pathology in mice that cannot expand their CAG tracts somatically. Conversely, Inhibition of DSBs reverses neuronal toxicity in animals that undergo somatic expansion. Although they coexist in neurons, DSBs and somatic expansion are independent therapeutic targets for HD.

## INTRODUCTION

Huntington disease (HD) is one of a class of neurodegenerative diseases for which unstable expanded (>35) CAG is the mutation underlying disease progression and severity^1–2^. HD is fatal. Pathophysiology of HD is characterized by a slow and steady decline of neurons, leading to a gradual deterioration of motor and cognitive abilities over time^1–4^. Despite years of effort, there is no cure^2^. Small molecule approaches to offset the clinical features of HD have been extensive ^2,5–1^. ^8^. While treatments to suppress chorea have had some clinical success^9^, the effects on neuropathology have been limited ^5,6,10^. The CAG tract in the mutant protein codes for a long polyglutamine region in the expressed gene product^1–4^. Thus, it has made intuitive sense that lowering the level of the mutant huntingtin protein (mhtt) would be a good approach to offset disease^11,12^. Although promising in model organisms, gene-silencing of the mutant allele by interference (RNAi) and microRNA strategies^13–16^, antisense oligonucleotides (ASOs) systems^17–21^, and Clustered Regularly Interspaced Short Palindromic Repeats/Caspase 9 (CRISPR/Cas9) systems^22,23^ has not yet succeeded in rescue of the disease phenotypes in clinical trials^24–27^.

The strong inverse relationship between age-at-onset of motor signs and length of the CAG repeats, however, has pointed to the expansion itself as the primary determinant of disease onset ^28–30^. Indeed, in humans and in model systems for disease, the longer tracts have earlier onset and more severe HD phenotypes ^30–36^, which appear to have a greater dependence on the somatic tract rather than the encoded polyglutamine protein^37–40^. Indeed, CAG to CAA variants have been detected in some HD patient alleles ^38,39^ which shorten the uninterrupted CAG tract but have no impact on polyglutamine length. In humans, loss of the CAA interruptions is associated with earlier onset^38,39^. The dependence of toxicity on the uninterrupted allele length has been confirmed in mice^37,40^. The somatic CAG tract rather than the polyglutamine protein is apparently the rate driver for toxicity.

Given its importance, use of gene editing to shorten the somatic tract length has been the focus of significant therapeutic efforts^38,41–45^. The DNA repair machinery are strong modifiers of somatic expansion, with MMR proteins being perhaps the most important enhancers^41–45^. In GWAS, disease-accelerating loci in HD include *MLH1* (MutL homolog 1), *PMS1* (post meiotic segregation increased 1 homolog), *MSH2* and *MSH3* (MutS homologs which dimerize to form MutSβ), *PMS2* (post meiotic segregation increased 2) and *LIG1* (DNA Ligase 1)^44,45^. The remarkable convergence of the GWAS on the MMR components has validated years of study corroborating a role for them in promoting somatic expansion in models for disease^46,47^. In a recent analysis, for example, loss of all nine GWAS-implicated MMR proteins suppressed somatic expansion and delayed disease onset in an HD mouse model expressing 140 repeats^48^. A CRISPR knockout screen in Htt^Q111^ mice^49^ identified MMR proteins and gap filling polymerases as strong enhancers of somatic expansion. Additionally, there was also a strong correlation in the CRISPR screen among repair genes that reduce CAG tract length with delay of HD onset. For example, Fanconi anemia nuclease 1 (FAN1), which is strong modifier associated with delay in human HD^50–5^ is a strong repressor of CAG expansion^49,51^. *MSH3(-/-)* or *FAN1(-/-)* have similar effects in mice expressing CGG^53^ and GAA^54^ repeats, indicating that the impact of DNA repair proteins on tract length is relevant in other expansion-related diseases. Relationships between the MMR systems and FAN1 are emerging^55–57^.

Strategies to reduce somatic CAG expansion are shaping new approaches to develop urgently needed therapeutics for HD. Despite its central importance, however, emerging evidence suggests that somatic CAG expansion, although necessary for disease, may not be sufficient for neuronal toxicity. In human HD tissue, for example, CAG expansions are prominent in medium spiny neurons (MSNs) of the striatum (STR) and in cholinergic interneurons and cerebellar Purkinje cells (PC), but only striatal projection neurons are lost, even though all these cell types express MSH3 and huntingtin proteins^58^. Since cell types can differ in their somatic CAG length, the differential vulnerability may, in part, be explained by threshold effects. A recent RNA Seq and modeling analysis indicates that striatal projection neurons in HD patients become vulnerable to death only after expansion reaches a somatic threshold of 150-180 CAGs, which occurs over 5-6 decades^59^. Early transcriptional changes, however, become prominent in the cell types which are most susceptible to death and may contribute significantly to neuronal loss^59^. Indeed, *zQ175/MSH3(-/-)* mice expressing a huntingtin allele with 175 polyglutamine repeats but lacking MSH3 cannot expand the CAG tract somatically, yet these animals develop transcriptional dysfunction as early pathology^60^. Thus, there are likely to be additional DNA dependent processes that contribute to HD toxicity and may be influenced by DNA repair.

We have investigated the expression and activity of five major DNA repair pathways for their ability to influence HD toxicity in mice. We report here that DSBs drive HD toxicity by mechanisms that are independent of somatic expansion. While somatic expansion is driven by active MMR, DSBs are driven by a loss of DSBR activity, which promotes transcriptional pathology and neuronal death. Although they occur together, CAG expansion and DSBs make independent contributions to toxicity. Therapeutic suppression of both may be critical in treating HD.

## RESULTS

### CAG expansion is progressive in the brains of *HdhQ(150/150*) mice

We have extensively evaluated and previously published the results of toxicity studies in *HdhQ(150/150*) mice^8,36,61,62^ . These animals harbor a long CAG tract of roughly 150 knocked into the endogenous mouse gene^63^ and expand it during life^34,36,63^. Motor abnormalities are detected around 20 weeks (wks)^8,63^ and neuronal death in the STR was detected around 60 weeks^8^, while pathology was spared in the CBL^8^ (Fig. 1A). The progressive pathology in *HdhQ (150/150*) mice is accompanied by CAG somatic expansion, as measured by us^8,36,61^ and others^34,35,63^(Fig. 1A). Somatic expansion in *HdhQ(150/150*) animals begins around 11 wks^34–36^ is prominent by 30 wks (Supplemental Fig. 1), and tract size is longer in the affected striatum (STR) relative to the resistant CBL^34^. Thus, CAG expansion in *HdhQ(150/150*) animal displays the genotype, age-and region-specific features of disease toxicity observed in humans.

**Fig. 1:**
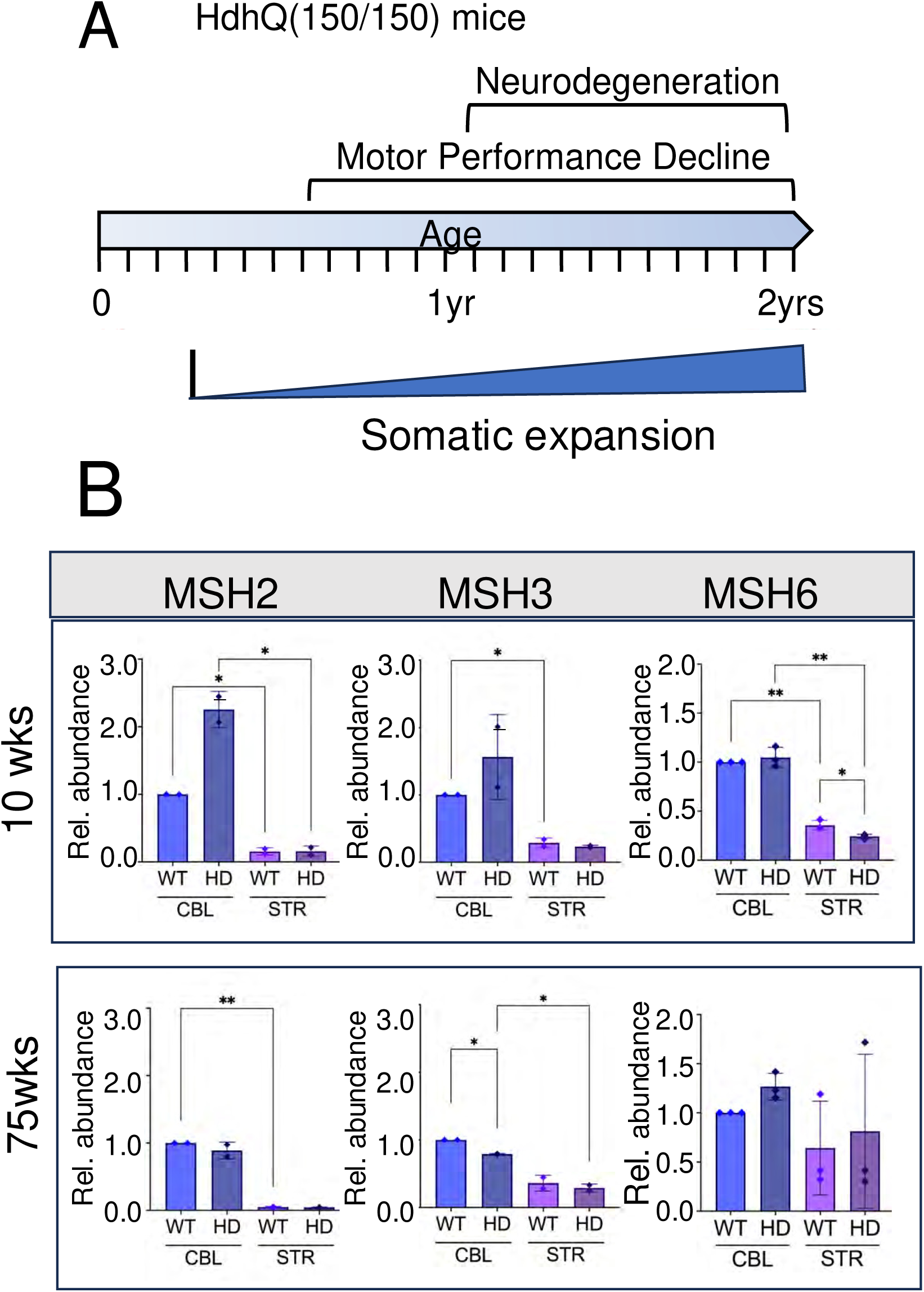
Mouse brain expresses the MMR recognition machinery necessary for expansion WT and *HdhQ(150/150*) mice. (A) Schematic diagram of timing and properties of pathophysiology and somatic expansion in *HdhQ(150/150)* animals over 2 years, summarized from^8^. Representative examples of Genescan for repeat sizing at 30 weeks are provided in Supplemental Fig. 1A. (B) Western analysis of MSH2, MSH3, and MSH6 in the affected STR and the resistant CBL in n=3 animals of 10-11 wks and three n=3 animals between 74-76 wks (referred to as 10 wks and 75wks) from brain extracts of *HdhQ(150/150)* animals and age and gender matched *C57BL/6J* controls (referred to as HD and WT, respectively). Two samples were resolved side by side indicated by the number 1 and 2 (Supplemental Fig. 2A) and data for a third animal is presented in the source file data. Six technical replicates of the SDS-PAGE gels are shown of each genotype at 10wks and 75wks. (Supplemental Fig. 2A). Each replicate gel was transferred to membranes and probed with specific antibodies to the indicated protein (P) or to GAPDH (C) with the molecular weight markers (kD) to the side of each gel (Supplemental Fig. 2A) The relative abundance of MSH2, MSH3 and MSH6 expression was normalized relative to the WT CBL. Protein antibodies are listed in Supplementary Table 1. Error bars represent standard deviation with the minimum (lower bar) and maximum (upper bar) values for the samples at each age. Significance was determined by a 1-way ANOVA; P is * 0.01 < p ≤ 0.01, **0.001 < p ≤ 0.01.

CAG expansion depends on the dose of MSH3^64–66^. In other lines, mice that expressed the highest MSH3 level expanded the most^66^. Thus, we expected that MSH3 expression in *HdhQ(150/150*) mice would track with the somatic expansion and display its properties, i.e., MSH3 expression would be highest in the STR. MSH2 dimerizes with MSH3 to form MutSβ and with MSH6 to form MutSα. However, MutSβ drives somatic expansion of CAG triplet repeats in knockout mouse crosses^67,68^, while MutSα has variable or random effects^67,68^. Thus, MSH2 served as a positive control for the effects of MSH3, while MSH6 served as a negative control in linking expression to somatic expansion. Clonal mouse strains express the same protein profile. Thus, we confirmed that *HdhQ(150/150*) mice express MSH2, MSH3 and MSH6 by measuring their levels in brain extracts from the sensitive STR and in the resistant CBL in six genetically identical *HdhQ(150/150*) mice, n=3 at young (10 wks.) and n=3 old (75 wks) ages relative to comparable WT controls (Fig. 1B). The resolved proteins from six replicate sets of SDS-PAGE gels were transferred to membranes and probed with specific protein antibodies to one of the MMR recognition proteins (P) or GAPDH (C). The plotted results of the protein expression analysis are shown (Fig. 1B), together with the accompanying gel images (Supplemental Fig. 2A). As judged by staining intensity, both WT and *HdhQ (150/150*) animals expressed MSH2, MSH3 and MSH6 in the STR and in the CBL of young (10 wks) and old (75 wks) mice (Fig. 1B). However, the expressed level of MSH3 and MSH2 was low in the affected STR of *HdhQ (150/150*) animals, which expanded the most, and, in contrast to the MSH6, did not increase with age as did progressive expansion.

Somatic expansion is most prominent in neurons^35,58,69,70^. Thus, it was possible that subcellular localization of MSH3 in *HdhQ (150/150*) animals changed with age and altered the regional properties of MSH3 expression in unexpected ways. To establish the cell type-specific properties of MSH3 expression, we measured it together with MSH6 at the singe cell level in brain tissue sections from WT and *HdhQ (150/150*) mice (Fig. 2). Protein expression was measured by immunofluorescence (IF) using specific antibodies and was assigned as neuronal if antibodies to the expressed protein co-stained with the NeuN(+) neuronal marker^71^ and, as glial, if the IF staining occurred in NeuN(-) cells in the tissue sections (Fig. 2A). Indeed, in magnified cell images from the tissue section in young and old animals, MSH3 (MSH3, green) co-stained with nuclear DAPI (Blue) and NeuN (purple) in both the STR (Fig. 2A, M, N, D) and CBL (Fig. 2B, M, N, D) of WT and *HdhQ(150/150)* animals, indicating that expression was highest in neurons in both regions. Additional IF images for the neuronal and glial expression of MSH3 in the STR of these animals are shown (Supplemental Fig. 2B, C). The cell type specific expression profiles for MSH3 (Fig. 2C, D) and MSH6 (Fig. 2E, F) were quantified in whole tissue sections as the average IF intensity from 50 neurons (NeuN(+)) cells and 50 glia (NeuN(-)) in randomly selected fields from n=3 animals. MSH3 was primarily neuronal in the STR of both genotypes (green (WT) and (red (HD)) and did not change with age (Fig. 2D). The MSH6 control was also expressed in striatal neurons and glia around 10-20wks (Fig. 2 E, F), but expression increased in neurons from *HdhQ (150/150)* mice relative to WT animals as these animals aged (Fig. 2F, 70-90wks). Collectively, the results confirmed that MSH3 was available to drive somatic expansion in neurons of both STR and CBL in *HdhQ(150/150)* mice, but the dose of MSH3 did not fully match the region-specific or age-dependent patterns of somatic expansion or HD toxicity in these animals.

**Fig. 2:**
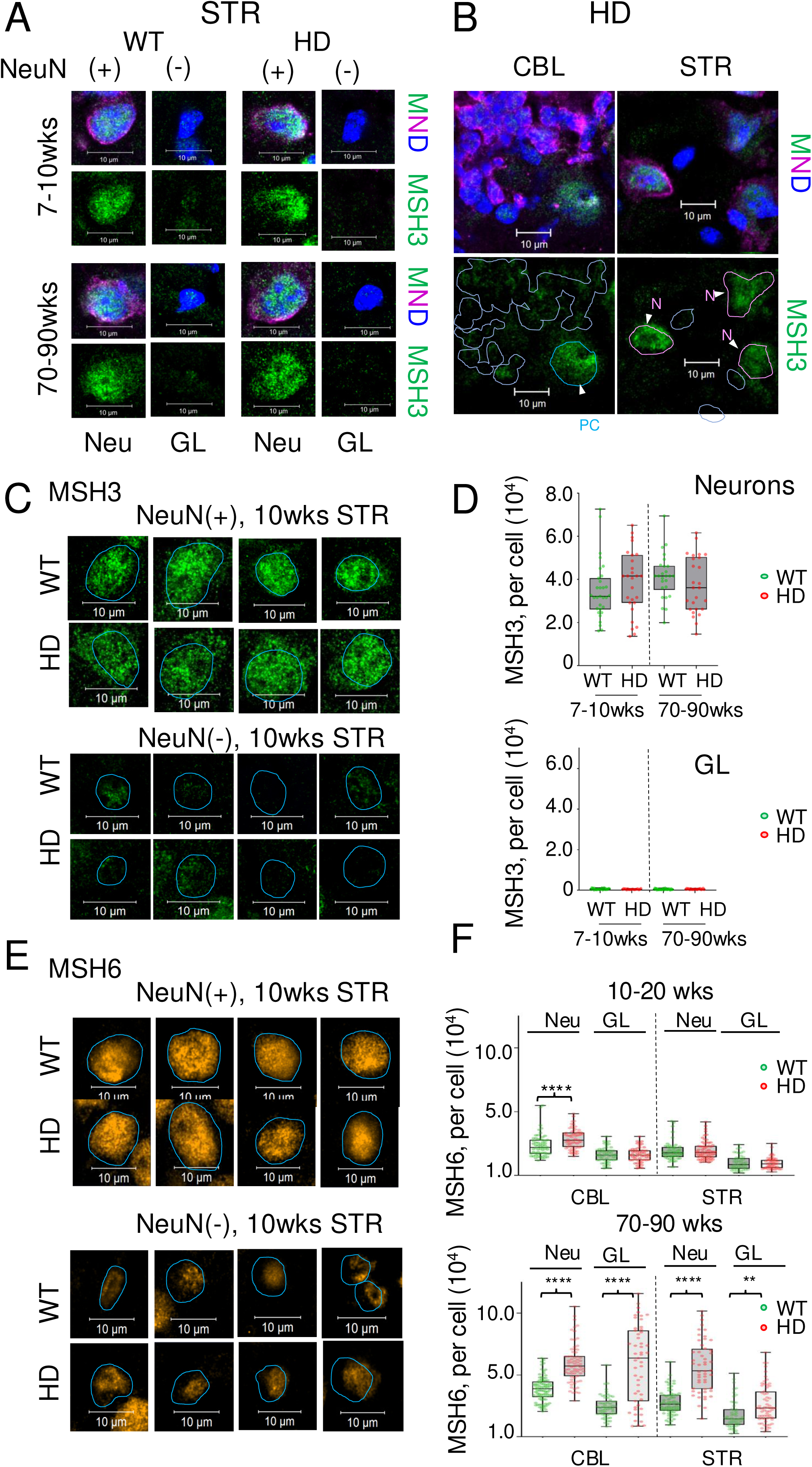
MSH3 and MSH6 expression is prominent in neurons of WT and *HdhQ(150/150*) mice. Single cell analysis of the MSH3 in the affected STR and resistant CBL of WT and HD animals. (A) Magnified images of MSH3 immunofluorescence (IF) for antibodies in individual cells within tissue sections from the STR of WT or HD animals of 15-20 wks or 70-90 wks, as labeled; (M, MSH3, green), (N, NeuN, magenta), (D, DAPI, blue). Scale bar is 10µm. The nuclear marker, DAPI (blue), neuronal marker, NeuN (N, purple) and MSH3 (M, green) staining are shown as an overlay (M,N,D) or MSH3 as an individual channel image (MSH3). NeuN(+) cells are neurons and NeuN(-/-) cells are glia. MSH3 co-stained prominently in NeuN(+) cells (purple) independent of genotype. More examples shown for STR of WT and HD animals in Supplemental Fig. 2B,C. (B) Same as (A) but comparing the IF of MSH3 expression in STR and CBL of 7-10 wk HD animals: (Left, CBL) The nuclear contours from the DAPI stain outlined in light blue indicate the position of the nucleus and highlight the poor MHS3 protein (green) staining intensity in most cells of the CBL; (Right, STR) MSH3 is strongly expressed in striatal neurons, which are delineated by the purple outline, and poorly expressed in DAPI(+), NeuN(-) glia. Scale bar is 10µm. (C) Magnified images of neurons (top) and glia (bottom) in the STR of 10wks WT and HD animals detected by IF antibody staining for MSH3. MSH3 expression is highest in neurons. (D) Quantification of MSH3 expression as determined by the average per cell staining intensity of 50 NeuN(+) neurons (top plot) and 50 NeuN(-) glia (bottom plot) in the striatal tissue sections from n=3 in WT(green bars) or HD (red bars) animals at 7-10wks or 70-90 wk. Data are displayed as a box and whisker plot, where the box are 25-75% of the values, the line indicates the median value, and 25% maximum values and 25% minimum values are indicated by whiskers above and below the box, respectively. (E) Same as C for MSH6 (orange), (F) same as E for MSH6. Significance was determined by a 1-way ANOVA; P is **0.001 < p ≤ 0.01 and **** p < 0.00001.

### Expression of DNA repair machinery is not significantly different in the affected STR and resistant CBL of WT and *HdhQ(150/150*) mice

Mechanisms for DNA repair are well characterized^72–74^. However, crosstalk has long been implicated between mismatch repair (MMR) machinery, double strand break repair (DSBR) ^75,76^, interstrand cross-link (X-linked pathway)^55–57^, transcription-coupled repair (TCR)/Nucleotide excision repair (TCR/NER)^77,78^, and base excision repair (BER)^79^. If a DNA repair pathway other than MMR was impaired or failed to remove DNA damage, then inefficient DNA repair might drive, at least in part, the patterns of toxicity in *HdhQ(150/150*) mice, independently of MSH3-driven somatic expansion. Thus, we tested (1) whether the machinery for other major DNA repair pathways was expressed (Fig. 3) and (2) whether they were active (Fig 4) in the brains of *HdhQ(150/150*) mice compared to WT controls.

**Fig. 3.**
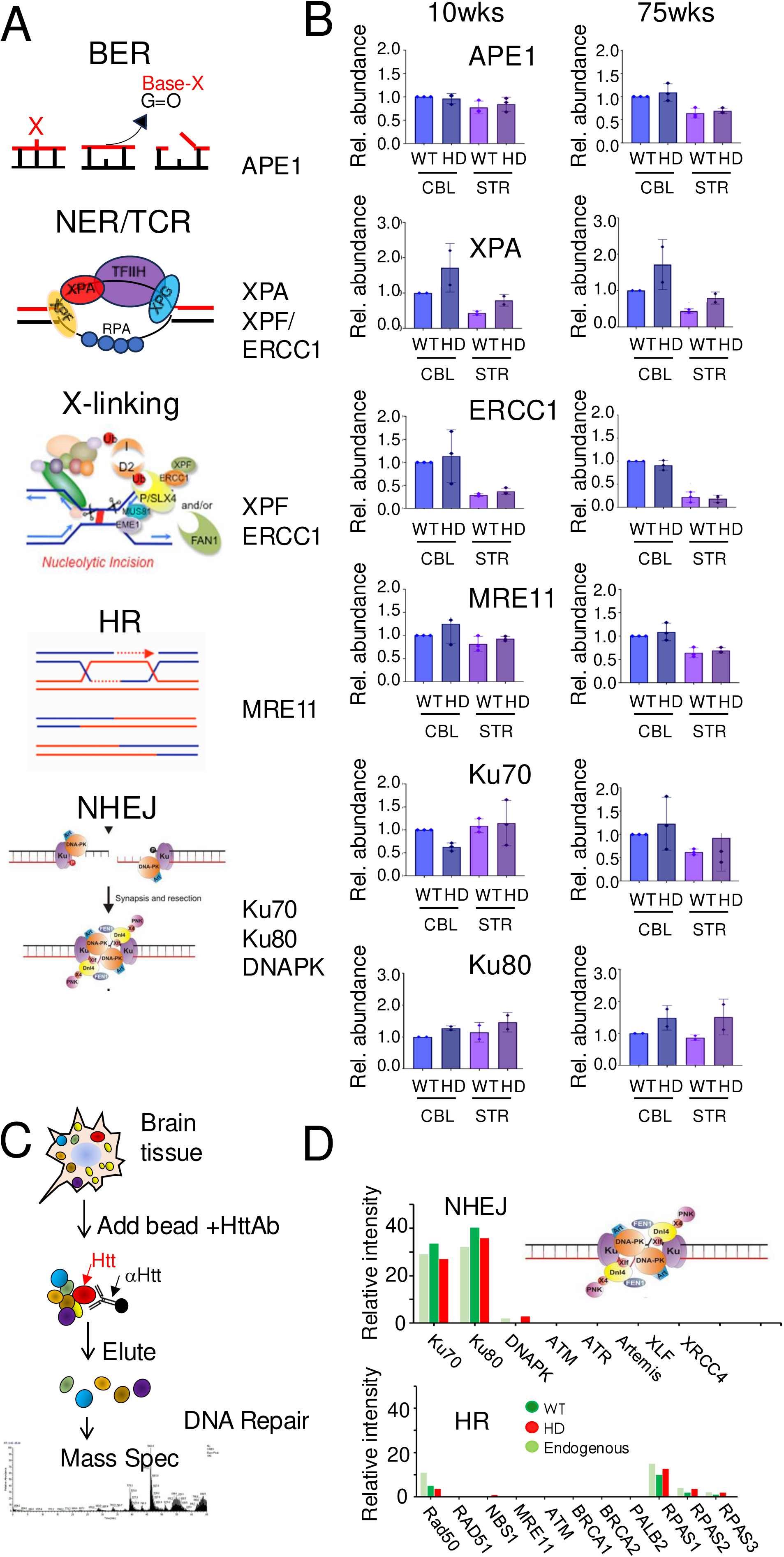
Htt and mhtt interacts with the DSBR machinery in tissues from WT and *HdhQ(150/150*) mice. (A) Schematic diagram of major DNA repair pathways^72–74^. The representative pathway proteins measured in (B) are indicated to the right of the schematic. (B) Western blot detection is the same as described in Fig. 1C but for Apurinic/apyrimidinic (AP) endonuclease (APE1); Xeroderma Pigmentosum Group A protein (XPA), Xeroderma Pigmentosum Group F protein (XPF), and Excision Repair Cross-complementation group 1 (ERCC1); Meiotic Recombination 11 Homolog 1 (MRE11); X-ray repair cross-complementing 6 (Ku70); X-ray repair cross-complementing 5 (Ku80), at the indicated ages. The proteins extracts were resolved in ten technical replicate SDS- PAGE gels; five gels for proteins measured in 10 wk animals (left) and five for proteins measured in 75 wk animals (right). Each blot was probed with indicated antibodies to a representative pathway protein (P) or GAPDH (C) (Supplemental Fig. 3). The protein expression data are normalized relative to the WT CBL. Error bars represent standard deviation with minimum (lower bar) and maximum (upper bar) values for the clonal samples. Source data and Full uncropped gels are provided in Supplementary Source File. The protein antibodies are listed in Supplementary Table 1. (C,D) Immunoprecipitation/Mass spectrometry (IP-MS) analysis of the interactions of DNA repair proteins with htt or mhtt. (C) Schematic diagram of the IP-MS analyses. (D) The IP-MS spectrometry results for endogenous NIH3T293 cells (light green) or for NIH3T293 cells overexpressing htt (dark green) or mhtt (red). (Top) Only NHEJ components or (bottom) HR components were identified among the pathways. The analogous IP-MS analysis of striatal tissue from WT and *HdhQ(150/150*) mice is shown in Supplemental Fig. 4.

**Fig. 4:**
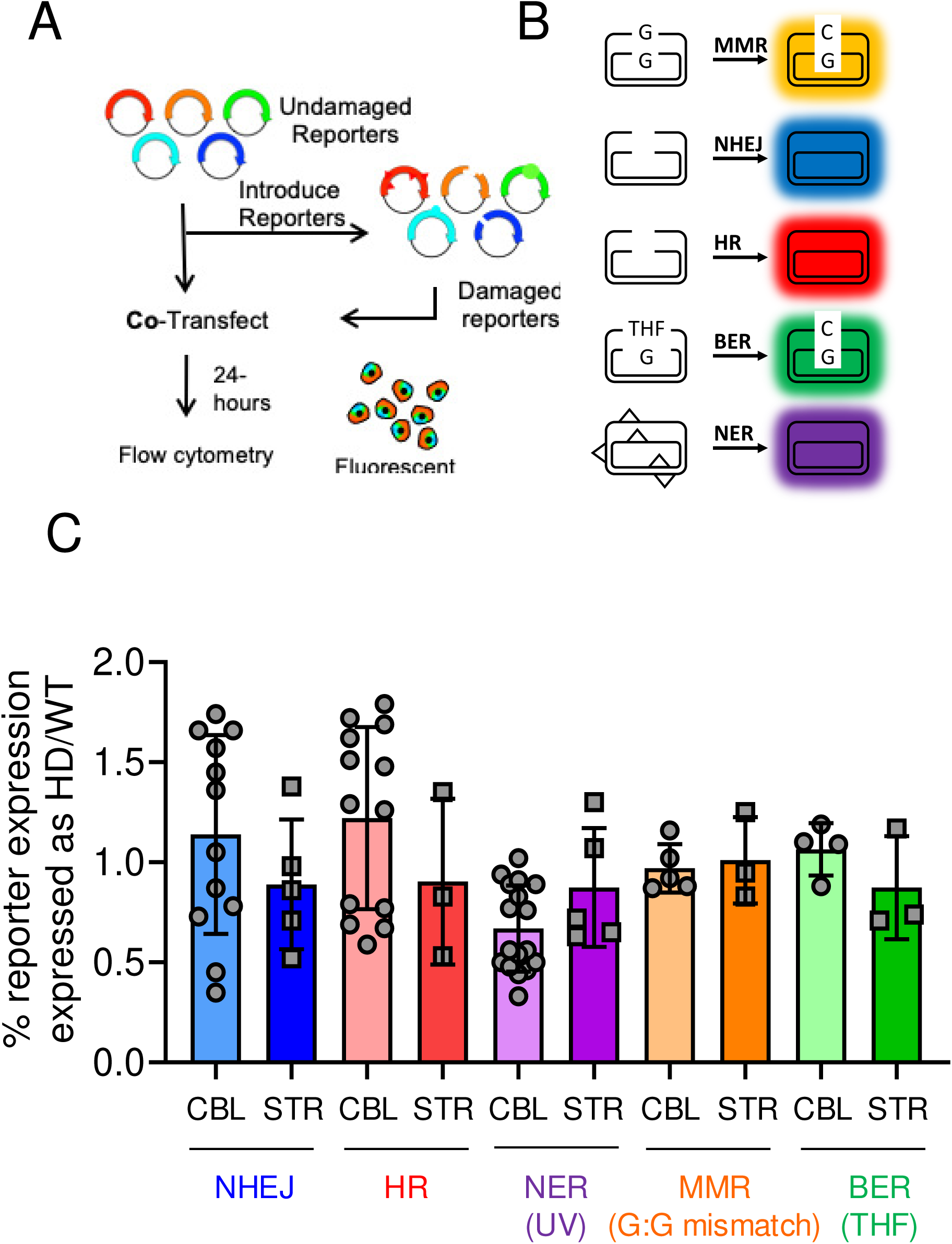
Mhtt inhibits the activity of NHEJ pathway of DSBR. (A) Schematic for the fluorescence multiplex host cell reactivation (FM-HCR) assay of different DNA repair pathways in primary glial cultures from the STR and the CBL of WT and HD animals. (B) Schematic diagram of the color-coded reporter plasmids for the indicated pathways; HR, NHEJ, MMR, NER, BER. Primary glia were seeded and transfected with sufficient efficiency (4- 8%) to afford a robust analysis of fluorescence intensity by flow cytometry^82–84^ and DNA repair capacity in the transiently transfected cells, described in text. (C) Activity plotted as % reporter expression of repair pathway in WT (light colors) and HD (darker colors) for each pathway, normalized for WT CBL. Repair activities were measured in plated glial cultures; n = 6 for CBL and n = 5-12 for the STR. Data are displayed as a box and whisker plot, where the box are 25- 75% of the values, the line indicates the median value, and 25% maximum values and 25% minimum values are indicated by whiskers above and below the box, respectively.

The protein expression was measured for the machinery needed to carry out common pathways of DNA repair including Homologous Recombination (HR), Non-Homologous End Joining (NHEJ), base excision repair (BER), nucleotide- or transcription coupled excision repair (NER/TCR), and DNA crosslink repair (X-link repair) (Fig. 3A)^72–74^. Due to the complex and multi-component nature of repair complexes, we did not evaluate all proteins in each pathway. Rather, we selected one or two representative proteins which were key to pathway function (indicated to the right in Fig. 3A). The protein expression for the excision pathways included the Apurinic/apyrimidinic Endonuclease 1 (APE1) of the BER pathway, Xeroderma Pigmentosum type A protein (XPA) for NER/TCR, and Xeroderma Pigmentosum type F (XPF) and type G (XPG) nucleases for TCR/NER (Fig. 3A). Although the Excision Repair 1 Endonuclease Non-Catalytic Subunit (*ERCC1*) and its binding partner XPF serve as the 5’ core nuclease for bubble excision in of NER/TCR, the dimer is also essential during steps of X-link repair (Fig. 3A). Proteins of the classic double strand break repair (DSBR) pathways included the meiotic recombination 11 (MRE11) (HR), and Ku70 and Ku80, proteins of NHEJ, which are encoded by the X-ray repair cross-complementing protein 6 and 5 genes, (*XRCC6)* and (XRCC5), respectively,

Brain extracts from the STR and CBL were prepared from four WT and four *HdhQ(150/150*) animals which were selected from the colony, two young animals (7-10wks) and two old animals (70-90wks) for each genotype. Ten replicate sets of SDS-PAGE gels (five for young and five for old ages) were transferred to membranes and probed with specific antibodies to a representative protein (P) from the pathways or with GAPDH (C). The potted results (Fig. 3B) and the gel images (Supplemental Fig. 3) are shown. As judged by antibody staining, brain regions of young (around 10 wks) and old (around 75 wks) animals of both genotypes expressed the representative proteins from all five DNA repair pathways tested (Fig. 3B). Toxicity associated with *HdhQ(150/150*) animals did not appear to correlate with alterations in the expression level of common DNA repair pathway components (Fig. 3A).

### The normal and mutant huntingtin protein interact with the machinery of NHEJ and HR

Since brain cells were equally well equipped with the machinery to repair DNA damage, the genotype-specific nature of toxicity raised the possibility that the mutant huntingtin (mhtt) might interact with the DNA repair machinery in *HdhQ(150/150*) animals differently than in normal huntingtin (htt). To test this idea, we used a specific htt antibody to immunoprecipitate htt-binding partners and tested whether DNA repair proteins were among the pull down products identified by mass spectrometry (IP-MS) (Fig. 3C and Supplemental Fig. 4A). In the first experiments, we evaluated the protein in tissues from the affected STR of WT and *HdhQ(150/150*) animals (Supplemental Fig. 4B). As judged by IP, htt in extracts from WT tissue and *HdhQ(150/150*) animals interacted with DSBR machinery. Indeed, the htt antibody “pulled down” Ku70 and Ku80, the core dimer for the end joining activity of NHEJ, together with RPA170, RPA32 and RPA14, which serve as general single strand annealing machinery, in both WT and *HdhQ (150/150*) animals (Supplemental Fig. 4B). The interactions were specific since no pulldown products were observed when a non-specific control antibody was used in the IP reaction (Supplemental Fig. 4C). While both Htt and mhtt interacted with the NHEJ machinery, neither protein associated with components of BER, NER/TCR or MMR pathways.

To detect whether there were weaker interactions, we repeated the tissue experiment in immortalized human NIH393T cells before and after overexpressing htt or mhtt (Fig. 3C, D). Like the tissue results, the htt antibody pulled down Ku70 and Ku80, RPAS1-3, in endogenous NIH393T cells, together with a minor amount of DNAPK catalytic subunit from the same pathway (Fig. 3D, light green), establishing NHEJ as a major target. The results confirmed the specificity of the mhtt-Ku70/Ku80 interaction since the same product was recovered using either truncated^80,81^ or full-length huntingtin proteins (Fig. 3D). To test for weak huntingtin interactions, we overexpressed htt and mhtt in human NIH393T cells using CMV-driven huntingtin cDNA plasmids harboring 26 or 51 CAG repeats, respectively. However, the results were unchanged; the NHEJ and RPA components remained the major pull down products together with an additional weak association with RAD50 and NBS1 of the HR pathway. The same IP-MS products were observed whether htt (Fig. 3D, dark green) or mhtt protein was overexpressed (Fig. 3D, red). Thus, the NHEJ machinery interaction was not specific for the mutant form of the protein.

### mhtt inhibits activity of NHEJ and HR in animals

Given that toxicity in *HdhQ(150/150*) animals depends on genotype, the IP-MS results raised the possibility that the interaction with mhtt selectively impaired DSBR activity *in vivo.* Thus, we directly evaluated the activity of the DSBR machinery in cells from WT and *HdhQ (150/150*) animals, alongside the activity of unaffected DNA repair pathways as negative controls (Fig. 4). The collective activity profile of DNA repair was measured using fluorescent multiplex host cell reactivation (FM-HCR) assays^82–85^ in primary glia from WT and *HdhQ (150/150*) animals. Primary glial cultures express the same DNA repair proteins as do neurons but at lower levels and have served as a good assay system for DNA repair in normal brain cells^85^. Briefly, a series of fluorescent reporter plasmids were engineered with DNA lesions that are specifically repaired by one of the major DNA repair pathways (Fig. 4A). The presence of the lesion alters the coding sequence of a fluorophore, preventing fluorescence intensity of the expressed reporter gene. Fluorescence intensity is restored when the reporter lesions are repaired (Fig. 4B). The reporter lesions included an overhang or blunt double strand breaks (DSB) for HR and NHEJ, respectively, a tetrahydrofuran abasic site analog (THF) for long patch BER, an A:C mismatch for MMR, or low ultraviolet (UV) radiation-induced DNA damage for NER (Fig. 4B). The efficiency of repair is calculated from fluorescence intensity of the reporter as a percentage of the expression from an equivalent reporter in the absence of damage. The normalization and calculation of repair capacity from flow cytometric data has been described in detail previously^82^. A major strength of this approach is the ability to measure the activity of DSBR simultaneously with activity of the other DNA repair pathways in the same cell. We refer to the collective pathway activity as a “DNA repair landscape”. If mhtt specifically impaired DSBR pathways, we expected that their activity alone would be affected in mhtt-expressing cells.

As measured by FM-HCR, all DNA repair pathways in the landscapes of WT and *HdhQ (150/150*) animals were active to varying degrees in cultured glial cells from STR of either genotype (Fig. 4C, normalized to the WT CBL). However, the activity of neither BER, MMR, nor TCR/NER depended on genotype (Fig. 4C). DSBR pathways, however, were inhibited in glia from HD animals. The activity of both HR and NHEJ activity is naturally low in the WT brain^85^, but it was further reduced in primary glia of *HdhQ (150/150*) animals (Fig. 4C). Thus, the FM-HCR assay supported the results of the pull-down experiments; only components of DSBR pathways had measurable interactions with mhtt (Fig. 3C, D), and only those were inhibited by mhtt in activity assays (Fig. 4C).

We validated the reduction in DSBR activity, in parallel, by quantifying the removal of DSBs induced after exposure to radiation (2Gy). The production of DSBs was detected by the appearance of foci from DSB marker γH2AX (Fig. 5A), and repair efficiency was monitored by the loss of γH2AX foci over a 24 hr post-irradiation interval (Fig. 5B, C). Low levels of radiation did not kill cells (Supplemental Fig. 5), but elevation of oxidative DNA damage can lead to DSBs when radiolysis of water generates hydroxyl radicals (OH)^86^. If HD brain cells had a functional deficit in repairing DSBs, we expected that, after exposure to radiation, loss of the γH2AX foci in the primary glia would be slower in cells isolated from *HdhQ(150/150*) mice relative WT cells. Indeed, radiation damage resulted in a prominent formation of γH2AX foci formation in glia from both WT and *HdhQ(150/150*) mice, confirming that radiation induced DSBs (Fig. 5A). In glial cultured from the CBL, loss of γH2AX foci was rapid and equivalent in both WT and *HdhQ(150/150*) cells (Fig, 5B, left). In the STR of *HdhQ (150/150*) mice, however, loss of γH2AX foci was substantially inhibited, and removal was slow compared to WT mice during the 24 hr observation period (Fig. 5C).

**Fig. 5.**
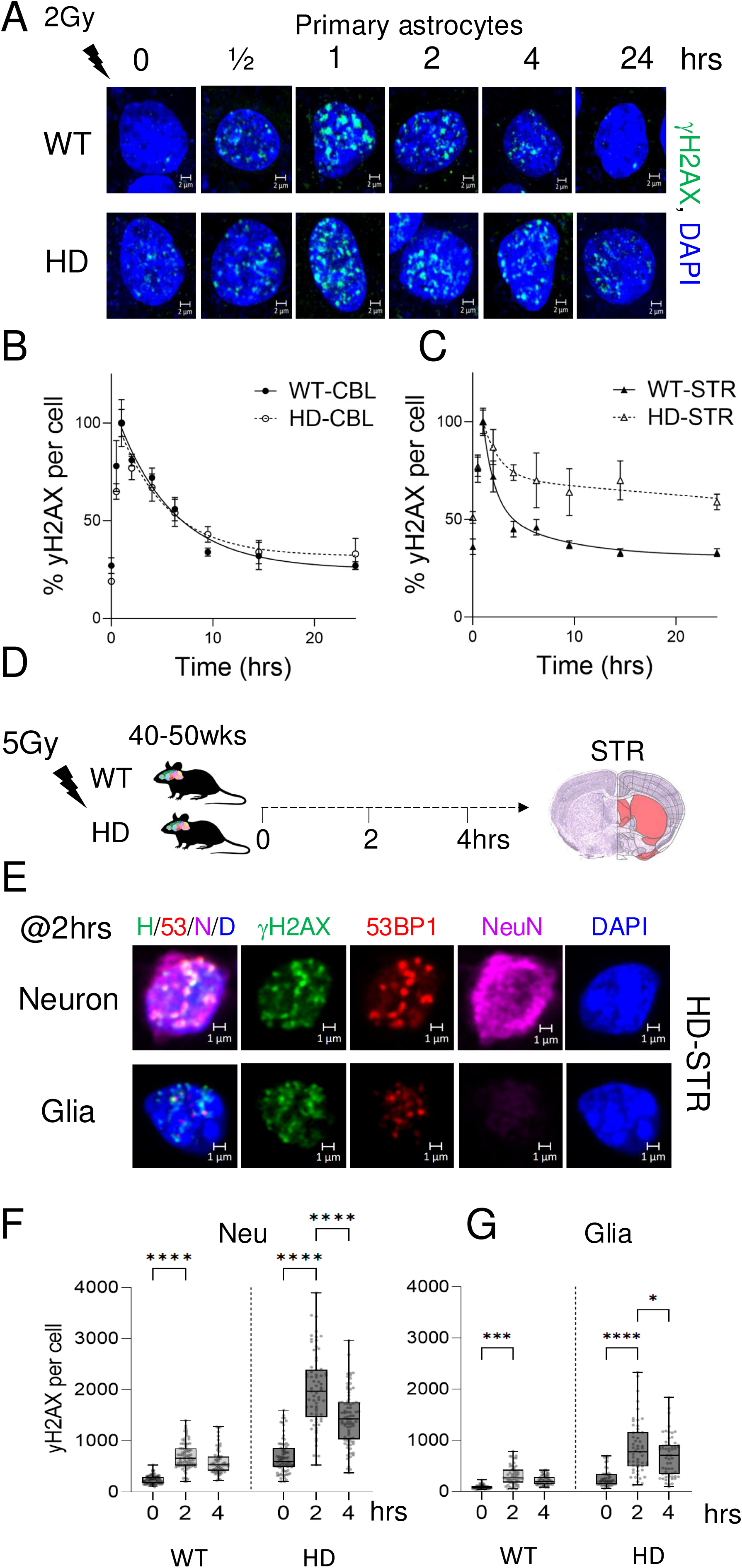
Repair of Induced DSBs is inhibited *in vitro* and *in vivo*. (A) The dynamics of γH2AX foci in primary glia cultures from STR in WT (top) or HD (bottom) animals exposed to 2Gy irradiation over 24hrs. The cells were co-stained with γH2AX (green) and DAPI (blue); γH2AX foci are turquoise in the overlay images. Scale bars is 2µm. (B) γH2AX foci signal intensity plotted as a function of time (hrs) and quantified in at least 100 glia cells from the CBL(left) and STR (right) of WT and HD animals in cultures from n=3 dissections. (D) Schematic illustration of DSB induction by radiation damage in the STR of WT and HD animals of 40-50wks. Animals were irradiated with a single 5Gy dose at the beginning of the experiment and the generated DSBs were quantified by γH2AX foci formed up to 4 hours post irradiation. Allen brain Atlas image indicating in red the STR tissue tested^117^. (E) Examples of striatal DSBs 2 hours post- irradiation in magnified images of neurons and glia of WT or HD animals at 70-90wks. Shown is IF intensity from co-staining with γH2AX (green), NeuN (purple), DAPI (blue), or 53BP1 (red), as separate channel images (panels 2-5) or as an overlay of all four stains (H/53/N/D, panel 1) in striatal tissue sections from HD animals. Co-staining of γH2AX and NeuN(+) cells defined DSB localization in neurons, while γH2AX in NeuN(-) cells were taken as DSBs in glia. (F,G) Quantification of DSBs at the single cells level for neurons (F) and glia (G) in whole tissue sections from WT or HD animals up to 4 hours post-irradiation, as judged by the staining intensity of γH2AX. Points shown are individual scored γH2AX signals from at least n=50 cells per tissue section (n=3 animals). Data are displayed as a box and whisker plot, where the line indicates the median value, the box are 25-75% of the values, and 25% maximum values and 25% minimum values are indicated by whiskers above and below the box, respectively. The probability statistics for all comparisons was determined from a 1-way ANOVA. P is * 0.01 < p ≤ 0.01, **** p < 0.00001.

### Induced DSBs *in HdhQ(150/150*) mice are inefficiently repaired *in vivo*

We tested whether Inhibition of DSBR occurred *in vivo* (Fig. 5D-G). The radiation experiment was repeated in living WT and *HdhQ(150/150*) mice by exposing adult animals (40-50wk) to a higher radiation dose (5Gy) (Fig. 5D). The animals were sacrificed immediately after irradiation or at 2 or 4 hours post irradiation (Fig. 5D, n=3 for each group). The level of DSBs were measured in the tissue sections from the STR of WT and *HdhQ(150/150*) mice using IF intensity of the γH2AX or 53BP1 DSB markers (Fig. 5E-G). In these experiments, the brain sections were stained with NeuN to identify neurons, and co-stained with nuclear DAPI (blue), γH2AX (green) and 53BP1 (red) to identify if neurons or glia were preferentially susceptible to DSBs (Fig. 5E). A representative result is shown (Fig. 5E). Each antibody stain is displayed as an individual color channel in the magnified cell image (Fig. 5E, panels 2-5) or as or an overlay of all four stains/antibodies (Fig. 5E, H/53/N/D, panel 1). Indeed, antibodies to both the γH2AX and the 53BP1 preferentially co-stained with NeuN(+) neurons of *HdhQ(150/150*) mice (Fig. 5E, top panels), while staining was weaker in NeuN(-) glia (Fig. 5E, bottom panels). Thus, DSBs formed most efficiently in neurons, suggesting that the damage was cell type specific. The repair of DSBs was quantified by the average IF staining intensity of γH2AX (green) in 50 NeuN(+) cells and 50 NeuN(-) cells randomly selected over the tissue field during the four-hour period. As judged by the weak γH2AX staining intensity, the DSBs in the STR were efficiently repaired during the observation period in both neurons (Fig. 5F, WT, Neu) and glia (Fig. 5G, WT, glia) of WT animals and DSBs in the glia of *HdhQ (150/150*) animals (Fig 5G, HD glia). However, γH2AX staining intensity significantly increased in striatal neurons of *HdhQ(150/150*) mice (HD) relative to WT mice (roughly 4-10-fold)(Fig. 5F, Neu), indicating that DSBR was inhibited in the affected brain region (Fig. 5F, Neu). Thus, DSBR in *HdhQ(150/150*) animals was suppressed in the STR of *HdhQ(150/150*) animals *in vivo* and *in vitro*. Suppression of DSBR was genotype, cell type- and region-specific.

### DSBs accumulate in the brains of *HdhQ(150/150*) mice together with transcriptional pathology

If inhibition of DSBR contributed to HD toxicity in mice, then we expected that DSBs would be elevated beyond normal levels in neurons of *HdhQ(150/150*) mice relative to WT animals. Furthermore, we expected that DSBs would accumulate to the highest extent with age in the STR of the disease animals. Indeed, in the STR (Fig. 6A) and CBL (Fig. 6B) of whole brain sections (Fig. 6C,D), the nuclear and peri-nuclear periphery of neurons in *HdhQ(150/150*) mice co-stained with the antibody to ATM-dependent KRAB-associated protein 1 phosphorylation (pKAP-1)^87^, which acts with γH2AX in recruitment of nucleosome remodeling activities. pKAP staining was prominent in the STR of *HdhQ(150/150*) mice (Fig. 6C, HD) relative to WT mice (Fig. 6C, WT) and increased with age from 7 to 100 weeks. Interestingly, in human patients, CAG expansion occurs in cerebellar PCs although the CBL is more resistant to cell death^58^. In the CBL of *HdhQ(150/150*) animals (Fig. 6D, HD), pKAP staining intensity was largely restricted to a single cell layer of large cells, which lay adjacent to dense neuronal layer (Fig. 6D and Fig. 6E, panel 1), and stained with a calbindin marker for PCs (Fig. 6E, panels 2-4)^88^. Staining by calbindin and pKAP was accompanied by significant autofluorescence (Fig. 6E, green) from oxidized lipofuscin lipids which has been reported previously^8,61^. Thus, in *HdhQ(150/150*) animals, striatal neurons, which were preferentially targeted for toxicity, accumulated pKAP with age, while staining was infrequent in most resistant cerebellar cells (Fig. 6D).

**Fig. 6:**
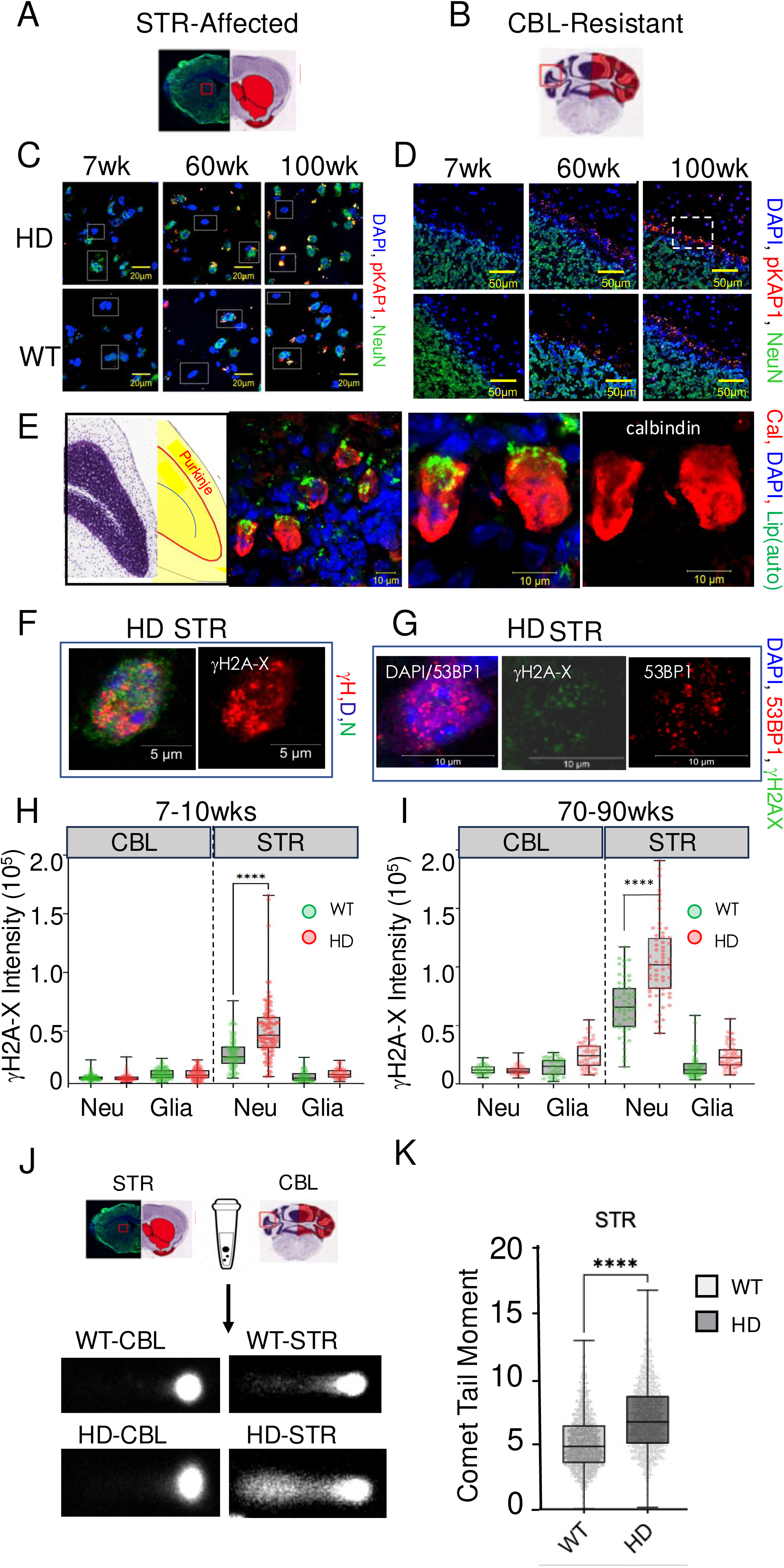
DSBs accumulate in neurons of HD mice. (A,B) Allen brain Atlas image^117^ for STR(A) and CBL(B) in H&E-stained experimental tissue sections. NeuN staining of the section is shown to the left in the ST. The segments used for imaging are indicated in red. (C) Representative IF images of striatal tissue sections for HD (top) compared to WT (bottom) mice, stained with antibody to pKAP (red), nuclear DAPI (blue), and NeuN (green) of male mice at 7, 60, and 100wks. Scale bar is 20µm. (D) Same as (C) for CBL. Little pKAP staining is observed in cerebellum except for a single Purkinje cell (PC) cell layer adjacent to dense granular neurons. Scale bar is 50µm. (E) Diagram of PC layer in the mouse CBL (panel 1). (Panel 2) A 5-fold magnified image of the PC layer in the white hatched box in D (100wk) stained with DAPI (blue) and calbindin (red), a marker for PC cells. Scaler bar is 10µm. Green emission is generated from the lipid autofluorescence^8^ (panel 3,4). A two-fold magnification of PC in panel 2, with (panel 3) or without (panel 4) the lipofuscin autofluorescence signal. (F) (left) Overlay image of co-staining with DAPI (blue), NeuN (green) and γH2AX (red) in a magnified image of a striatal tissue section from an HD mouse of 70-90 wks; (right) γH2AX (red) alone. Scale bar is 5µm. (G) Same as (F) co-stained with two DSB markers in the same cell: (left) overlay image of DAPI (blue) and 53BP1 (red); (middle) γH2AX (green) alone; (right) 53BP1 (red) alone in a separate channel image. Scale bar is 10µm. (H,I) Single cell quantification of DSBs in neurons and glia from the IF intensity in tissue section from WT and HD mice at 7,8 wks (H) and 70-90wks. (I). Neurons were identified as NeuN(+) cells in the tissue section (see F). WT is green, HD is red. γH2AX staining was quantified from n=50 randomly selected cells of each type in n=3 tissue sections of 7,8wk (left) or 70-90wk (right) animals, respectively. Data are displayed as a box and whisker plot, where the box are 25-75% of the values, the line indicates the median value, and 25% maximum values and 25% minimum values are indicated by whiskers above and below the box, respectively. The probability statistics for comparing significance among regions were determined from a 1-way ANOVA are **** P < 0.0001 for the neurons. (J) Examples of comet tails from dispersed cells collected from the STR (top, left) and CBL (top right) of WT and *HdhQ(150/150*) tissue sections, as indicated. The imaging software delineates the comet head and the tail portion, which is visible for the STR. (K) Quantification of comet tail moments for the STR of WT (light gray) or HD (dark gray) mice at 70-90 wks. Points shown are individual scored comets from at least n=550-890 cells per tissue section (n=3 animals). Data are displayed as a box and whisker plot, where the box is 50% of the values, the line is the median value, and 25% maximum and 25% minimum values are indicated by whiskers above and below the box, respectively. Regional comparions were determined for signifiance using a 1-way ANOVA. ****, P < 0.00001 for the comparison.

To establish that pKAP staining reflected DSBs, we repeated the IF experiment in contiguous brain tissue sections of *HdhQ(150/150*) mice using antibodies to two additional DSB markers: γH2AX^89^ and 53BP1^90,91^ (Fig. 6F-K)(Supplemental Fig. 6). γH2AX forms a specialized chromatin structure that can extend hundreds of kilobases away from the DSB, while 53BP1 promotes repair of DSBs by NHEJ as part of the Shieldin complex^91^ and suppresses HR. Indeed, DSBs were detected primarily in neurons by the co-staining cells with γH2AX (red) and NeuN (green)(Fig. 6F) (additional examples in Supplemental Fig. 6). Furthermore, cells with a signal for γH2AX (green) also co-stained positively with the DSB marker 53BP1 (red) (Fig. 6G)(Supplemental Fig. 6). DSBs were quantified in the tissue of WT and *HdhQ(150/150*) animals from the average IF saining intensity of 53BP1 from 50 neurons (NeuN(+)) cells and 50 glia (NeuN(-)) cells in randomly selected fields in the STR and in the CBL from n=3 animals (Fig. 6H,I). As quantified by staining, DSBs were minimal in the CBL of either WT or *HdhQ(150/150* animals at any age (Fig. 6H,I, CBL). However, DSBs increased significantly with age in NeuN(+) cells in the STR from *HdhQ(150/150*) (red) mice relative to WT (green) tissue (Fig. 6H,I, STR, right). Thus, DSBs preferentially formed in striatal neurons and accumulated with age. DSBs mirrored the genotype-dependent, region-specific, cell type-specific, and age-dependent features of HD toxicity.

To confirm that staining with γH2AX and 53BP1 antibodies reflected actual DSBs, they were measured directly using a neutral comet assay^92,93^ at young (7wks) and old (75wks) ages (Fig. 6J,K). The details of the CometCHIP experimental parameters have been previously reported in the normal brain and the same methods were used^85^. Dispersed cells were collected and subjected to electrophoresis under neutral conditions (Fig. 6J) and the trailing DNA or “tails” were quantified by system software (Fig. 6K). Comet tails were low in the CBL (Fig. 6J). Consistent with the single cell analysis of γH2AX staining (Fig. 6I), however, comet tails were elevated in the affected STR of HD animals relative to WT (Fig. 6J, n=300-500 tails). Thus, DSBs co-existed with somatic expansion in the brains of *HdhQ(150/150*) mice. DSBs were present at the onset of somatic expansion (12 weeks) and preceded the early motor abnormalities (20wks) and neurodegeneration (60wks) in *HdhQ(150/150*) mice (see Fig. 1A).

### DSBs promote neuronal death independent of somatic CAG expansion

Since they occurred together with CAG expansion in *HdhQ(150/150*) mice, it was not possible to fully determine whether DSBs contributed to toxicity in this line. Thus, we used the *zQ175 /MSH3(-/-)* mouse as a separation of function mutant to segregate the impacts of the two types of damage (Fig. 7). The zQ175 parents harbor a CAG tract of 175 and exhibit significant HD-like phenotypes and seizures^94^. Mice that express a mutant htt allele but lack MSH3 cannot expand their CAG repeats somatically, and, indeed, *zQ175/MSH3(-/-)* crosses did not expand their HD alleles^60^. However, *zQ175* and the *zQ175/MSH3(-/-)* crosses, both of which expressed the Q175 protein, developed transcriptional dysfunction and protein aggregation as early pathologies (summarized in Fig. 7A)^60^. Thus, somatic expansion did not drive the development of early pathologies in *zQ175/MSH3(-/-)* mice.

**Fig. 7:**
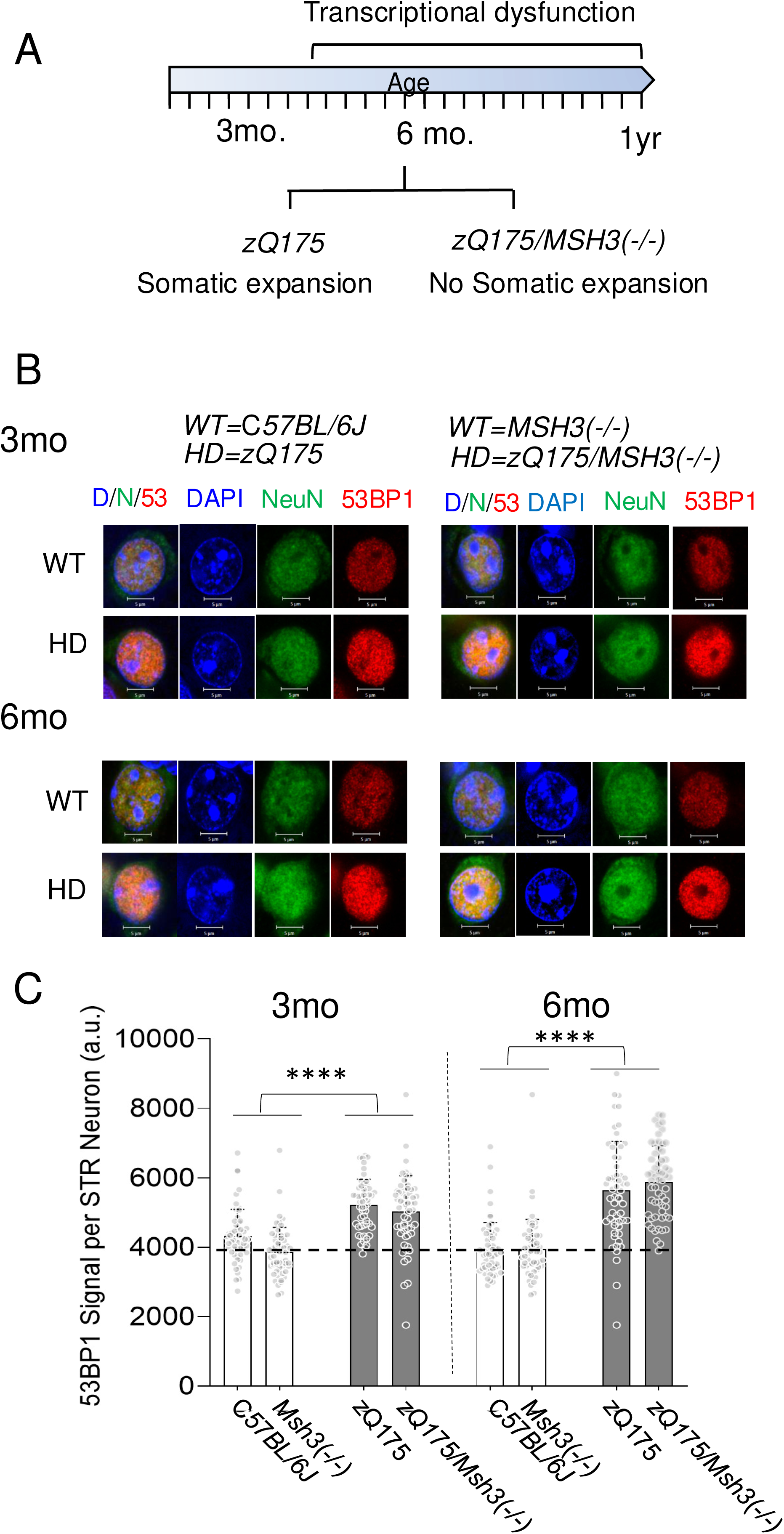
DSBs increase together with progressive transcriptional dysfunction in *zQ175/MSH3(-/-)* animals that cannot expand their CAG tract. (A) Summary of measured pathology and somatic expansion quantified in *zQ175* and in *zQ175/MSH3(-/-)* animals as published previously^60^. Somatic expansion is attenuated in *zQ175/MSH3(-/-)* animals. (B) The DSBs compared among tissue sections from WT, *zQ175/MSH(+/+), MSH3(-/-)* and *zQ175/MSH3(-/-)* at 3 and 6 months. Example of magnified neurons at 3mo (top) and 6mo (bottom) from brain tissue sections of (left) WT and *zQ175* mice and (right) *MSH3(-/-)* and *zQ175/MSH3(-/-)* mice stained with nuclear DAPI (blue)(panel 2), NeuN neuronal marker (panel 3, green), the 53BP1 DSB marker (panel 4, red) or an overlay of all three (D/N/53, panel 1). The genotypes designated as WT and HD are indicated at top. (C) Plot of DSBs as measured by 53BP1 staining intensity in neurons per indicated genotype at 3 or 6 months. At least n=50 cells per region were scored for DSBs in minimally three sections (n = 3). DSBs are elevated in *zQ175 and zQ175/MSH3(-/-)* animals at 3 and 6 months (gray bars), compared to the brains of WT or *MSH3(-/-)* control strains (white bars), respectively. Data are displayed as a standard deviation. Genotype comparions were determined for signifiance using a 1-way ANOVA. ****, P < 0.00001 for all comparisons.

To test whether DSBs contributed to pathology, the brain tissue from *zQ175* and the *zQ175/MSH3(-/-)* with transcriptional dysfunction was evaluated for DSBs compared to C*57BL/6J (WT)* and MSH3(-/-) controls (Fig. 7). Although the *zQ175* parental strain did not lose neurons at young ages between 3 and 6 months (Supplementary Fig, 7), DSBs were detected by IF staining with DSB markers in whole tissue sections in the brains of *zQ175* and the *zQ175/MSH3(-/-)* animals (Fig. 7B). Examples of tissue staining are shown (Fig. 7B). In a magnified cells from the tissue fields, some DSBs were measurable in all four mouse genotypes at any age, as detected using either γH2AX or the 53BP1 DSB marker probes (Fig.7B, compare 3 months and 6 months). However, the intensity of γH2AX and 53BP1 staining was higher in mhtt expressing strains, *zQ175* (left) and *zQ175/MSH3(-/-)*) (right), relative to the C*57BL/6J* (left) and *MSH3(-/-)* (right) controls strains, respectively, at both 3 (upper panels) or 6 (lower panels) months (Fig, 7B).

The DSBs in neurons were quantified by the average IF staining intensity of 53BP1 staining in minimally 50 NeuN (+) cells randomly selected over the tissue field for each genotype at 3 and 6 months (Fig, 7C). The 53BP1 signal in zQ175 and *zQ175/MSH3(-/-)* crosses were compared to those of the C*57BL/6J (WT)* and *MSH3(-/-)* parental controls, respectively (Fig. see 7B). If they were an independent driver of transcriptional dysfunction, we expected that DSBs would accumulate with age in *zQ175 and zQ175/MSH3(-/-)* crosses, relative to either WT animals or to *MSH3(-/-)* control animals. Moreover, the elevation of DSBs would accompany the progression of pathology to the same extent in *zQ175* and *zQ175/MSH3(-/-)* crosses. Indeed, DSBs were low in both WT or in *MSH3(-/-)* parental controls at 3 or 6 months (Fig. 7C, white bars). In contrast, a modest rise in the level of DSBs were obvious in both the *zQ175 and zQ175/MSH3(-/-)* genotypes by 3 months and continued to increase over 6 months, as judged by 53BP1 marker intensity (Fig. 7C. black bars). Indeed, in a short span of 3 months, DSBs increased in *zQ175 and zQ175/MSH3(-/-)* animals and were elevated by 30-50% relative to *WT* and *MSH3(-/-)* controls at comparable ages. Thus, DSBs accompanied the progression of early transcriptional dysfunction observed in these animals^60^. Although no somatic expansion was present in *zQ175/MSH3(-/-)* crosses, DSBs accumulated in this genotype to the same extent as in the *zQ175HD* parental mice. Collectively, the results indicated that MSH3 drove the formation of somatic expansion, but did not drive the formation of DSBs. DSBs in *zQ175/MSH3(-/-)* accompanied the development of early transcriptional dysfunction in the absence of somatic expansion.

### Correction of DSBs by XJB-5-131 in *HdhQ(150/150*) animals rescues toxicity with minimal effects on somatic expansion

If DSBs were a separable process contributing to HD toxicity, then reducing DSB formation should be beneficial, even in the presence of CAG expansion. We have previously measured DSBs in the CBL and STR of normal mouse brain (Fig. 8A, Allen brain atlas maps) before and after treatment with a mitochondrially-directed antioxidant, XJB-5-131 (Fig. 8B). Treatment suppresses DSB formation *in vivo* in normal control mouse brain^85^, in cultured primary glia^52,85^ and in NIH3T3 cells^85^. In *HdhQ(150/150*) animals, XJB-5-131 (Fig. 8C) prevented the onset of motor abnormalities (observed 20wks), prevented the decline in neurons (observed at 60 wks) if treatment began before onset of pathology^8,61,62^, and blocked disease progression if XJB-5-131 treatment was started after pathology developed^8^ (Fig. 8C). To determine whether DSBs drove the reversible pathology, we measured it together with somatic expansion and DSBs in the same tissues^8^ (Fig. 8D-G). Treatment with XJB-5-131 had only small effects on somatic expansion in the STR of these animals (Supplemental Fig. 8, shift from birth to 90wks ± XJB-5-131). However, XJB-5-131 suppressed DSBs as measured by the IF staining intensity of γH2AX (Fig. 8D). As shown in a representative magnified IF image of a striatal tissue section, γH2AX staining intensity increased in Vh-treated *HdhQ(150/150*) (HD,Vh) (Fig. 8D) compared to comparable WT animals (WT, Vh), and staining intensity was suppressed by XJB-5-131 treatment (Fig. 8D, HD, +XJB). In the same tissues, XJB-5-131 (+XJB-5-131) blocked the decline of striatal neurons that were lost in Vh-treated *HdhQ(150/150*) mice after 30 weeks of treatment (Fig. 8E)^8^. Neuronal toxicity was reversible by suppressing DSBs, without significant effects on somatic expansion.

**Fig. 8:**
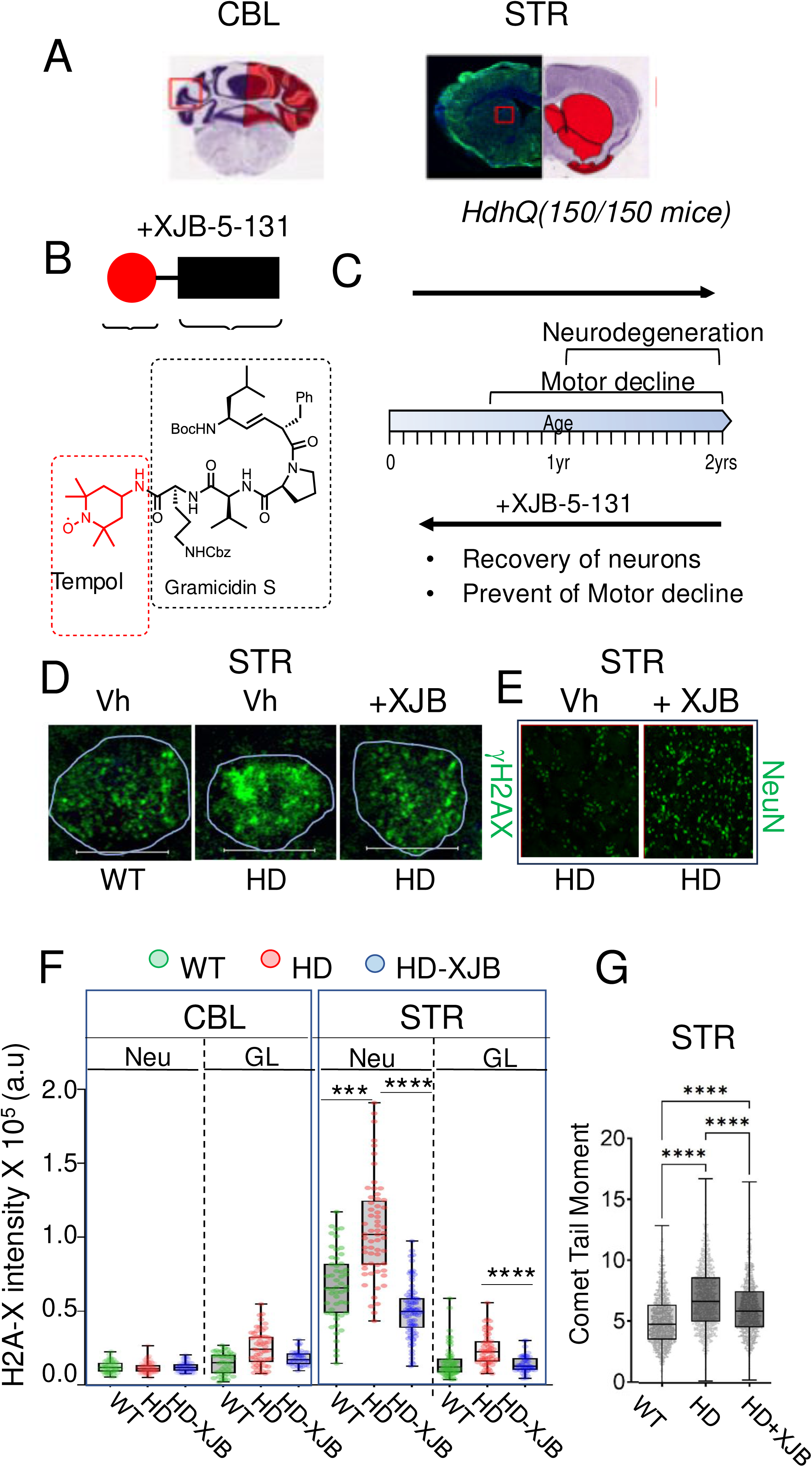
XJB-5-131 treatment of *HdhQ(150/150*) mice attenuates DSBs and prevents neuronal death. (A) Allen Brain Atlas map of affected STR and the resistant CBL for reference^117^. (B) Diagram of functional residues of the XJB-5-31 ROS inhibitor and its chemical structure. The red ball represents the tempol antioxidant (red) (red hatched box), which is fused to a mitochondria-targeted carrier peptide (black) (black hatched box). The target carrier peptide is based on the Gramicidin S, an antibiotic that targets the mitochondrial membrane directly. XJB-5-131 prevents base oxidation and SSB to DSB conversion. (C) WT or HD animals were aged to 60wks to allow accumulation of DSBs, before starting treated with saline vehicle (Vh) or XJB-5- 131 for 30 wks. Brain sections from 90wks animals were harvested and the sections were evaluated for DSBs, neuronal loss in tissue sections, and somatic expansion in dissected tissue from a contiguous section. Summary of reversible pathology quantified in *HdhQ(150/150*) mice before and after XJB-5-131 treatment^8^. (D) DSBs detected in brain sections by the IF signal intensity of the γH2AX DSB antibody marker (green) in WT or *HdhQ(150/150*) mice. Vehicle (Vh) was saline. Shown are magnified striatal neurons from Vh-treated WT (WT, panel 1), Vh-treated *HdhQ(150/150*) mice (HD, panel 2), and XJB-5-131 treated *HdhQ(150/150*) mouse (HD +XJB, panel 3). Scale bar is 10µm. (E) Corresponding NeuN antibody staining for (D) in tissue fields for Vh (left) or XJB-5-131-treated (right) *HdhQ(150/150*)(HD) mice. (F) Single cell analysis of derived from γH2AX IF signal intensity of WT (*C57BL/6J*) mice treated with Vh (green); *HdhQ(150/150*)(HD) mice treated with Vh (red) or XJB-5-131 (blue). The quantification is the average of 50 randomly selected NeuN positive neurons (Neu) and 50 randomly selected NeuN(-) glia (GL) from CBL and STR of (n=6) mice of each treatment group. Data are displayed as a box and whisker plot, where the box is 50% of the values, the line is the median value with 25% maximum and 25% minimum values indicated by whiskers above and below the box, respectively. The significance of the DSB marker intensity between treatment groups were evaluated using a 1-way ANOVA, **** P < 0.00001. G. Neutral comet for DSBs. n=1300-1600 comet tails per sample. Dispersed brain cells from each sample were collected at 70wks for neutral Comet Assay for WT(Vh), (HD) (Vh) or HD(XJB-5-131) (dark gray), as indicated. Data are displayed as a box and whisker plot as described for F; the significance of the treatment specific comparisons of DSB marker level were evaluated using a 1-way ANOVA, **** P < 0.00001.

The DSB level was quantified in the same tissue sections by the average IF staining intensity of γH2AX (green) in 50 NeuN(+) neurons cells and 50 NeuN(-) glia in the STR and in the CBL of both genotypes and both treatment groups (Fig. 8F). DSB formation, as detected by γH2AX staining intensity, was minimal in both neurons (Neu) and glia (GL) in the resistant CBL of either WT (green) or *HdhQ(150/150*) (red) animals (Fig. 8F, CBL) or in glia from the affected STR of *HdhQ(150/150*) animals (Fig. 8F, STR, GL) independent of the treatment group. Under these conditions, XJB-5-131 had modest but measurable suppressive effects on the formation of DSBs in the STR (Fig. 8F, HD-XJB, blue). However, DSBs doubled in striatal neurons of Vh-treated *HdhQ(150/150*) animals (Fig. 8F, STR, red) relative to comparable Vh-treated WT controls (Fig. 8F, STR, green), and XJB-5-131 treatment blocked the rise (Fig. 8F, blue). To ensure that γH2AX staining reflected actual DSBs, we measured them directly by neutral Comet assay in dispersed cells from the same tissues (Fig. 8G). Indeed, the comet tail moments in the STR of *HdhQ(150/150*) animals (Fig. 8G) reflected the same XJB-5-13-reversible pattern as did γH2AX staining (Fig. 8F, STR, Neu). Thus, a progressive rise in DSBs accompanied the onset and drove the reversible pathology in *HdhQ(150/150)* mice. XJB-5-131 suppressed DSBs and mediated improvement in striatal pathology regardless of somatic expansion (Supplemental Fig. 8). The reduction in DSBs in the STR after XJB-5-11 treatment was directly proportional to the improvement of NeuN staining and motor performance in *HdhQ(150/150*) mice^8^.

## DISCUSSION

GWAS analysis has provided strong evidence that modifiers of CAG tract length have a crucial influence on HD onset^29,38,43–46,49^. Here, we report that DSBs are a second driver of disease pathology (Fig. 9), which does not depend on somatic expansion. DSBs and CAG somatic expansion co-exist in the HD brain, but the two types of DNA damage drive disease toxicity by distinct mechanisms. Active MMR promotes the site-specific increases in CAG tract length within the HD allele. Somatic expansion can proceed independently of DSBs if MSH3 and its associated factors are expressed. DSBs, on the other hand, do not depend on MSH3 (Fig, 7), and are driven by a loss of DSBR activity (Fig. 4 and 5). We propose a model in which CAG expansion and DSBs provide two mechanistically distinct drivers of HD toxicity. The CAG tract encodes an mhtt protein that inhibits NHEJ (Fig. 4), leading to elevated DSBs, which alter transcription and ultimately kill neurons. In the past, the small effects on CAG expansion by loss of DSBR proteins in mice argued against them being a strong factor in expansion associated diseases. However, DSBs do not impart their toxicity by changing the CAG tract length. Three key features define a specific role of DSBs as a driver of disease toxicity in *HdhQ(150/150*) mice with or without somatic expansion.

**Fig. 9:**
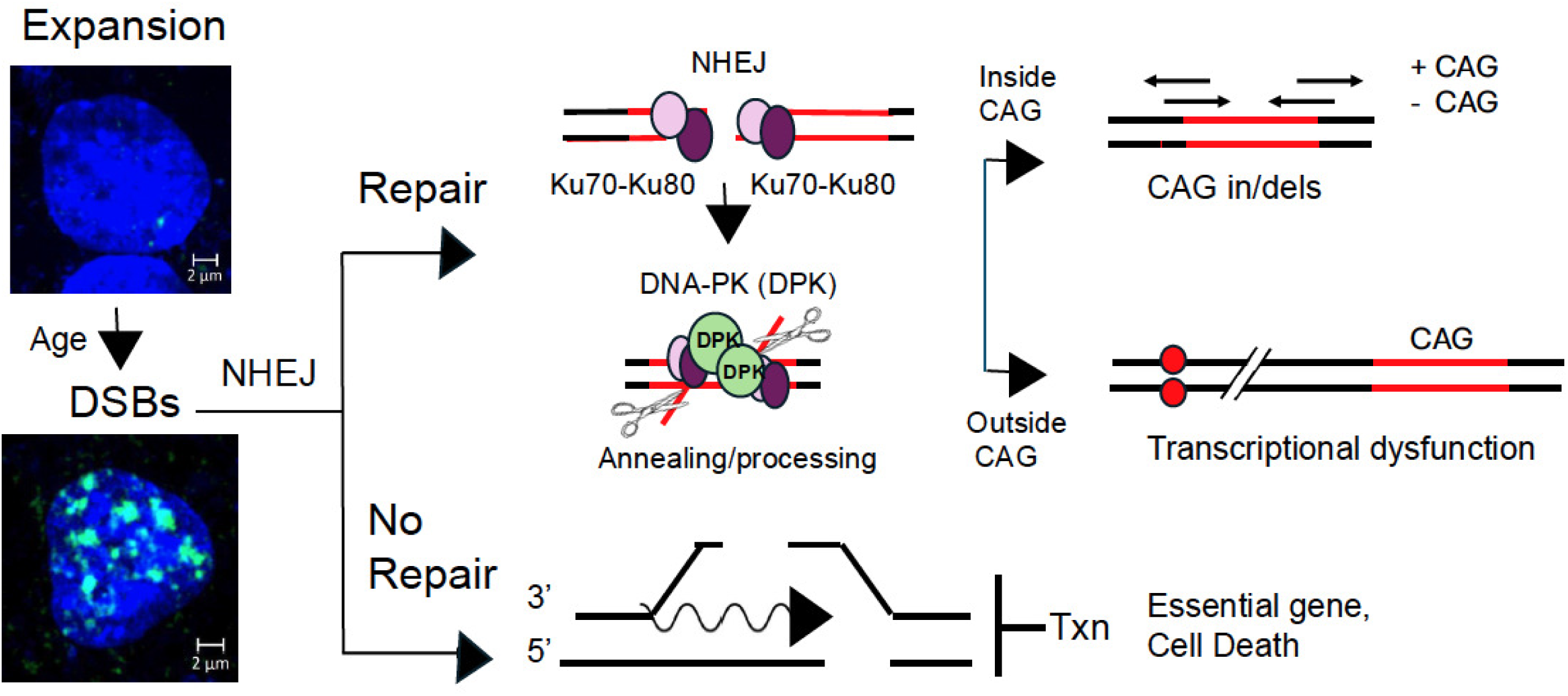
Model for the impact of DSBs in promoting HD toxicity. (Left) Cells with an expanded CAG tract (upper) develop DSBs with age (lower) as an indicated by accumulation of gH2AX foci staining. Scale bar is 2µm. (Right) Model for the role of DSBs in promoting HD toxicity. (Repair, NHEJ). NHEJ uses Ku70/Ku80 dimer to recruit DNAPK and the nuclease Artemis (scissors) to prepare two broken DNA ends for ligation. Extensive DNA end resection is prevented by Ku binding and artemis typically removes short regions of DNA to expose patches of microhomology. (Inside CAG tract) if the DSBs occur within a CAG repeat tract (red), microhomology is high and nuclease processing results in no change, or small insertions/deletions of CAG repeats. (Outside CAG tract). DSBs occur preferentially outside the CAG tract in HD animals. Annealing is imperfect at random genomic sequences. Artemis typically removes short regions of DNA to expose patches of microhomology and removes compatible bases to improve homology^95.97^, often resulting in somatic mutations in the genome (red balls). The resulting somatic mutation occur most frequently in gene enhancers. influencing transcriptional regulation^104^. (No repair) Unrepaired DSBs within a gene will terminate transcription, and if the DSB occurs in an essential gene, will cause death.

First, DSBs do not substantially change the length of the CAG repeat. NHEJ introduces repair errors when two strands around the break are pulled together to find homology for annealing and ligation of the broken ends^95,96^.(Fig. 9, Repair). Within the CAG repeats (Fig. 9, red lines), microhomology is high and annealing may be accurate. Although CAG repeats can fold back or generate overhangs, the artemis nuclease (Fig. 9, scissors) removes DNA ends at single-strand/double-strand junctions^96–98^, including structures that may arise between the two DNA ends being joined^98^. Since Ku70-Ku80 blocks removal of large DNA segments^96–98^, NHEJ results in only small CAG insertions or deletions, even if DSBs form within the CAG tract (Fig. 9, repair inside CAG, in/dels). Contractions are more likely than large expansion if overhangs are incompatible. These features provide a basis for the outcome in mouse models of losing DSBR machinery. Indeed, CTG repeats in myotonic dystrophy type 1 (DM1) mice lacking DNAPK catalytic subunit^99^ or CAG repeats in HD animals lacking Ku70 or Ku80^80^ display only small changes in tract size. Recent CRISPR knock out experiments in Htt^111^ mice reveal that NHEJ-associated ligase IV is a strong suppressor of CAG tract length^49^. HR does not compensate for inhibited NHEJ since there is no homologous chromosome in non-dividing neurons^95^. Indeed, CGG repeat expansion in a fragile X model is independent of DSBR-mediated by Pol θ, RAD52, RAD54 or RAD54B^100^, and loss of Rad 52 or Rad54 has little effect on somatic CTG expansion in the brains of myotonic dystrophy type 1 (DM1) mice^99^. Thus, DSBs and somatic CAG expansion are separable processes. Since NHEJ is inhibited, however, a DSB within the CAG tract may not be repaired, terminating transcription of htt, which is essential for life.

Second, most DSBs form outside the CAG tract in random genomic DNA (Fig. 9, black lines), and repair errors introduced during NHEJ alter gene expression. In the striatum of *HdhQ(150/150*) mice, we find DSBs are on the order of 10^5^, which is a residual population of those formed (Fig. 6), while the CAG repeat tract at the HD locus represents less than 10^-7^ of genome. Thus, some, if the not the majority of DSBs, will be located outside the CAG tract and will have no effect on its length. However, in random genomic DNA, homology for annealing and ligation of the broken ends is poor. Artemis (scissors) processing during NHEJ improves homology by removing incompatible sequences and mispaired bases, but at the expense of introducing changes in the DNA sequence at the repair junction (Fig. 9, repair outside CAG, red dots)^96–98^. Indeed, error-prone repair is a common source of somatic polymorphisms, which normally yield nucleotide variants throughout life^101–104^. Although the genomic locations of DSBs are not site-specific, clustering of genomic signatures confirms that error-prone DNA repair targets somatic mutations to active genes^104^ and influences their transcriptional activity^103–107^. For example, in the absence of a replication fork, it is generally accepted that most DNA damage in neurons arises from random endogenous base damage (alkylation, oxidation or deamination), which occurs tens of thousands of times per day^108–110^. Removal of chemically modified DNA bases leaves transient SSB intermediates, which can convert to a DSB if two lie simultaneously on opposite sides of the helix^85,108,109^. Repair-Seq^111^ or Synthesis-associated with repair sequencing (SAR-Seq)^112^ have been used to map the sites of breaks by the incorporation of biotinylated nucleotides during repair. Sequencing by either method confirms that SSBs and DSBs are closely clustered at high levels within the enhancers of mammalian genes^112,113^.

DSBs also arise during active transcription itself, from concurrent SSBs on the opposing DNA strands^108–110^. For example, if RNAP encounters an abasic product of BER or a structural impediment, DSBs can arise in the process of resolving the block when one SSBs is introduced by TOPI cleavage and the other at the resulting R-loop behind the polymerase^114,115^. In normal cells, it now recognized that DSBs have functional roles in gene expression^116^. Topoisomerase II (TOP2)-mediated DSBs stimulate the expression of early response genes (c-*Fos*, *FosB*, *Npas4*, and *Egr*1) ^117,118^, which serve as functional adaptors for transient modulation of beneficial downstream activity. Thus, the elevated DSBs in HD, if repaired by NHEJ, are poised to increase nucleotide changes within transcriptional regulatory regions. Whether nucleotide changes in the genomes of HD patients drive transcriptional dysfunction remains to be validated, but it is a prominent feature of disease in both HD patients^29,59,58^ and in mouse models^37^. Indeed, our results from “separation of function” mice (Fig. 7) demonstrate that DSBs accumulate together with progressive transcriptional dysfunction in *z175Q/MSH3(-/-)* animals that cannot expand their alleles somatically.

Third, because NHEJ is inhibited, it sometimes fails to correct DSBs (Fig. 9, No repair). In non-dividing cells, DSBs are tolerated if a gene is dispensable. As DSBs increase in *HdhQ(150/150*) animals (Fig. 6), however, there will be an increasing risk that unrepaired DSBs will terminate transcription of an essential gene, leading to death (Fig. 9, No repair). Indeed, the inability to correct DSBs is toxic to neurons in HD animal models. We show here that suppression of DSBs by XJB-5-131 rescues neuronal death and reverses toxicity in *HdhQ(150/150*) mice even if somatic expansion is active (Fig. 8). Collectively, CAG expansion and DSBs provide two coincident but mechanistically distinct drivers of HD toxicity. The CAG tract encodes an mhtt protein that inhibits NHEJ (Fig. 4), while the resulting DSBs alter transcription and ultimately kill neurons (Fig. 9). Whether early transcriptional dysfunction or death occurs depends on where the DSBs are formed and whether error-prone NHEJ is successful in removing them (Fig. 9).

Although DSBs appear to be independent drivers of toxicity, the machinery for DSBR has not been identified in GWAS analysis of HD patients. DSBs, however, form randomly at loci throughout the genome and will vary among patients. It is also not clear whether loss of activity can be detected as a classic modifier. Since NHEJ activity is suppressed, partial effects may not provide sufficiently robust statistical correlations among DSBR genes and disease onset in HD patients. Nonetheless, it is notable that Ligase IV of the NHEJ pathway been identified recently as a repressor of CAG expansion in a mouse CRISPR knockout screen^49^. Why other components of the pathway have failed to be identified as modifiers of repeat length will need further experimental clarification. Regardless, DSBs co-exist with CAG somatic expansion and together lead ultimately to neuronal death. However, each process has a distinct driver, a distinct mechanism, and a separable location-dependent impact on toxicity. As such, these two types of DNA damage are independent therapeutic targets. Our data suggest that suppressing CAG expansion and DSBs together is likely to be a beneficial therapeutic strategy for HD and may be necessary to offset disease.

## METHODS

Our research complies with relevant ethical regulations.

### Breeding and maintenance of *HdhQ(150/150)* and C*57BL/6J* controls

The Institutional Animal Care and Use Committee approved all procedures. Animals were treated under guidelines for the ethical treatment of animals and approved by IACUC protocol #274005 at Lawrence Berkeley Laboratory. All animal work was conducted according to national and international guidelines. Male C*57BL/6J* were used in every experiment and referred to as C*57BL/6J* or wildtype (WT) mice (Jackson Labs cat#000664). Unless otherwise stated, the male C*57BL/6J* were used as controls in all analysis, referred to as WT. At least 3 animals (n=3) were used in each tissue experiment.

### Brain Tissue Lysate Preparation

Dissected brain tissues were flash frozen in liquid nitrogen and were preserved at -80°C until lysate preparation. Brain tissue was placed in a microcentrifuge tube and thawed on ice for approximately 10 min before adding lysis buffer consisting of T-PER Tissue Protein Extraction Reagent (Thermo Scientific, Cat No 78510), ∼8.7x Halt protease inhibitor cocktail (from 100x, Thermo Scientific, Cat No 78430), ∼4.3x Halt phosphatase inhibitor cocktail (from 100x, Thermo Scientific, Cat No 78420). After adding the lysis buffer, the tissue sat on ice for 2-3 min before being triturated with a Bel-Art extended handle pestle (Millipore-Sigma, Cat No BAF199210001). The triturated tissue sample was centrifuged at 16,100g, 4 °C for 1.5 min in Eppendorf 5425R refrigerated microcentrifuge. Following the centrifugation, the sample was subject to 4 cycles of pulse sonication on ice, performed in which 2 cycles of 10s-on and 30s-off followed by centrifugation at 16,100g, 4 °C for 1.5 min, 1 cycle of 10s-on and 30s-off followed by centrifugation at 16,100g, 4 °C for 1.5 min, and then 1 cycle of 10s-on and 30s-off. Sonication was performed using Branson Sonifier Cell Disruptor 185. The resulting homogenate was centrifuged once more at 16,100g, 4 °C for 30 min and supernatant was used for the protein quantification assay, SDS-PAGE and western blotting analyses.

### Protein Quantification Assay

The protein concentration of each lysate was quantified using Pierce 660 nm Protein Assay reagent (Thermo Scientific, Cat No 22660). Briefly, a small volume of each lysate was diluted 1:10 in phosphate-buffered saline (PBS). 10 μL of diluted lysate was mixed with 340 μL of Pierce 660 nm Protein Assay reagent in a 1.7 mL microcentrifuge tube and then incubated at room temperature (RT) for approximately 5 min before loading 150 μL of each mixture in each of 2 wells of a flat bottom, transparent 96-well plate. For each lysate, a mixture was prepared and analyzed in triplicate by Infinite M1000 microplate reader (Tecan) using Tecan i-control software.

### SDS-PAGE and Western Blot

SDS-PAGE samples were prepared by mixing brain tissue lysate containing 60 μg total protein (20 μg for apurinic/apyrimidinic endonuclease 1 or APE1, and 10 μg for glyceraldehyde 3- phosphate dehydrogenase or GAPDH), T-PER Tissue Protein Extraction Reagent (to level the volume of all the samples), 1x NuPAGE LDS Sample Buffer (from 4x, Invitrogen, Cat No NP0007) and 1x NuPAGE Sample Reducing Agent (from 10x, Invitrogen, Cat No NP0009). Each sample was boiled at ∼95 °C for 10 min and centrifuged at 15,000g, at RT for 5 min. Total proteins in each SDS-PAGE sample were subsequently resolved along a Novex WedgeWell Tris-Glycine 4-12 % Mini gel (Thermo Fisher, Cat No XP04205BOX) in XCell SureLock Mini-cell Electrophoresis system (Thermo Fisher). The resolved proteins were transferred to a nitrocellulose membrane using Trans-Blot Turbo Transfer System (Bio-Rad), with a standard protocol (25 V, 1.0 A, 30 min). The resulting nitrocellulose membrane was blocked in 5% non-fat dry milk in PBS + 0.05 % Tween-20 (PBST) for 1 hour at RT. The membrane was then incubated in 5 % non-fat dry milk in PBST + primary antibody for 1 hour at RT. The membrane was then washed 3 times with PBST. The membrane was subsequently incubated in 5 % non-fat dry milk in PBST + secondary antibody for 30 min at RT. The membrane was next washed 3 times with PBST and once with PBS. Lastly, Amersham ECL Select Western Blotting Reagent (Sigma-Aldrich, Cat No GERPN2235) was applied to the membrane and the western blot image was developed using VersaDoc MP 4000 Imaging System (Bio-Rad) and Quantity One 1-D Analysis software (Bio-Rad). Quantification of protein band intensity was performed using Image Lab software (Bio-Rad). The specific antibodies used in the analysis are listed in key resources Supplementary Table 2 and antibody testing is provided in Supplementary Table 3.

### Immunoprecipitation and Mass Spectrometry

Cell pellets or mouse brain tissue were minced and lysed in lysis buffer (IP Lysis buffer, Pierce #87787) containing protease inhibitor (Halt Protease Inhibitor Cocktail, Pierce #78429). Cell debris was removed using centrifugation (15,000g, 10min, at 4°C). Samples were immunoprecipitated with anti-Htt antibody (1:1:1 mix of Millipore#MAB2166, #MAB2168 and #MAB2170) bound to magnetic beads (Protein G Dynabeads, Invitrogen#10007D). Protein complexes were washed with 50mM TEAB (triethylamonium bicarbonate) buffer (x4) and digested off the beads using 0.0025% Trypsin (in TEAB) for 18hrs with shaking. Supernatant was collected and beads were washed with additional TEAB. (proteomic profiling) The combined eluants were analyzed using Data Independent Acquisition (DIA) and Liquid Chromatography tandem Mass Spectrometry (at the UCDavis Proteomics Core Facility). Expression plasmid used was the cDNA for the full-length huntingtin protein with 26Q (htt) or 51Q (mhtt) repeats, driven by the CMV promoter.

### Brain Tissue Sections

We collected brains from male mice (7-10 wks and 70-90 wks of age) of *C57BL6/J* mice. Brains were cut into 4 coronal sections and arranged in a holder filled with OCT (Tissue-Tek O.C.T. from Sakura), and immediately frozen in isopentane bath cooled by liquid nitrogen, prior to storage at -80°C. This arrangement of the tissue permitted concurrent cutting of all 4 sections at a time. Cuts were made so that all the relevant regions (caudoputamen of striatum, CA1 region of hippocampus, the granular and molecular layer of Crus1 of the cerebellum, and the entorhinal area of the cortex) were present in each cut. Sectioning onto slides (Histobond from VWR) was performed on a cryostat (Leica CM1950) using cut settings (chuck = -14°C, blade = -15°C) and cutting 10-15μm thick sections. Slides were air dried (15 min) and stored at -80°C until use. In parallel, the dissected tissue of equal weight before freezing was dispersed for cell number counting on a hemocytometer.

### Mouse brain fixation and sectioning of *zQ175 and zQ175/Msh3(-/−)* mice and controls

Tissue slices from the *MSH3(--), zQ175* and *zQ175/MSH3(-/-)* mice were provided from G Bates. The zQ175 Mice were transcardially perfusion fixed with 4% paraformaldehyde (Pioneer Research Chemical Ltd). Brains were removed and further post-fixed for 6 h at 4 °C before cryoprotection with gradient steps of 20% and 30% sucrose in 0.01 M PBS (Sigma). Once the brains reached equilibrium (had sunk to the bottom of the sucrose solution) they were washed in PBS, embedded in OCT (CellPath Ltd) and stored at −80°C. Coronal brain sections were cut at 30 μm on a cryostat and stored free floating at −20°C in tissue protective solution [30% ethylene glycol, 25% glycerol and 0.05% sodium azide in PBS] until staining.

### Immunofluorescence Staining and Imaging

Primary antibodies used were mouse anti-NeuN Alexa-488 conjugate (Millipore #MAB377X) (1:500), mouse anti-GFAP Cy3 conjugate (Abcam #ab49874) (1:500), mouse anti-APE1 (Novus #13B8E5C2) (1:500), mouse anti-Ku80 (Santa Cruz #515736) (1:500), mouse anti-ERCC1 (Santa Cruz #17809) (1:500), rabbit anti-MSH2 (Abcam #92473) (1:500), mouse anti-MSH3 (Millipore #MABE324) (1:500), rabbit anti-MSH6 (Abcam #ab92471) (1:500), rabbit anti-XPA (AbClonal #A1626) (1:500) and rabbit anti-MRE11 (Novus #NB100-142) (1:500). (Table S1) Secondary antibodies used were donkey anti-mouse Alexa-488 (Jackson #715-545-150), goat anti-mouse Alexa-568 (Invitrogen #A21124), donkey anti-rabbit Alexa-488 (Jackson #711-545- 152), and goat anti-rabbit Alexa-555 (Invitrogen #A32732). Anti-mouse antibodies were tested and selected for those having the least amount of background staining, which was typically visible as staining of blood vessels amongst various commercially available options.

Brain sections on slides were thawed and fixed with 4% PFA for 20min at 4°C, then washed once with PBS. They were then pre-extracted with RNase in CSK buffer, i.e., 0.3mg/mL RNase A (New England Biolabs) in (10mM PIPES pH 7.0, 100mM NaCl, 300mM sucrose, 3mM MgCl_2_, 0.7% Triton X-100). Lipofuscin autofluorescence was quenched by soaking in 1x TrueBlack (CellSignal #92401) in 70% EtOH (30sec) (care was taken not to allow sections to dry out), prior to 3 washes in PBS. Sections were then blocked with Fc Receptor Blocker (Innovex #NB309) for 15min at RT and then with Background Buster (Innovex #NB306-50) for 15min at RT, prior to washing once with PBS. Sections were then coated with 200μL of primary antibodies (1:500 diluted 1:500 in 10% Background Buster: PBS) and incubated for 1hr at 37°C or overnight at 4°C, prior to 2 washes of 10min each with PBS. Sections were then stained with 200μL of secondary antibodies (1:1,000 diluted in 10% Background Buster: PBS) and DAPI (10μg/mL) for 30min at 37°C. Finally, the slides were washed 2 times with PBS, 15min each, and refixed with 4% PFA for 10min. Sections were mounted using Immu-Mount (Epredia) and #1 coverslips (Electron Microscopy Services), sealed with nail polish, and stored at -20°C until they could be imaged.

### IF measurement of DSBs in tissues

Methods are the same as for IF detection of proteins except an antibody for γH2AX or 53BP1 was used (Supplemental Table 1). Tissue sections were co-stained with DAPI, an antibody to NeuN, and antibodies to γH2AX. DSBs in neurons were detected as co-staining of γH2AX and NeuN. Quantification of IF staining intensity for γH2AX was determined from 50 randomly selected cells of each type within the tissue sections or in cells.

### Brain sample preparation for the tissue Comet assays

Methods for Comet assays have been previously reported (85). Samples of young and old mouse brain regions (CBL, CTX, STR, HIP) were prepared before use for the Comet assay. Young (15 wks) and old (roughly 100 wks) C57BL/6 mice were sacrificed and 2-3 mm samples of the four brain regions of interest were collected, flash frozen and stored in liquid nitrogen until use. Samples were minced with scissors in 30 µL of buffer (HBSS, 20 mM EDTA, 10% DMSO) on ice. Next, 400 µL of buffer was added (200 µL for the HIP region, as it usually has the least cells). The mix was added on a 40 µm filter-top tube and centrifuged for 3 min at 150 g and 4°C. The cell concentration for each region was estimated by mixing 10 µL of each sample with 1 µL of 100x SYBR Gold (Invitrogen, Cat no. S11494), pipetting 10 µL on a hemocytometer (Hausser Scientific, PA, USA) secured on a glass slide and counting cells at 5X magnification using an inverted LED fluorescence motorized microscope (Zeiss LSM 710 microscope, Carl Zeiss Microscopy, GmbH, Germany). Every sample was brought to 300,000 cells/mL in 25 µL with PBS and used for the Comet assay.

### Brain cell preparations for CometChip assay

Samples of young and old mouse brain regions (CBL, CTX, STR, HIP) were prepared before use for the Comet assay. A young (15wk) and an old (100wk) C57BL/6 mouse were sacrificed and 2- 3 mm samples of the four brain regions of interest were collected, flash frozen and stored in liquid nitrogen until use. Samples were minced with scissors in 30 µL of buffer (HBSS, 20 mM EDTA, 10% DMSO) on ice. Next, 400 µL of buffer was added (200 µL for the HIP region, as it usually has the least cells). The mix was added on a 40µm filter-top tube and centrifuged for 3 min at 150 g and 4°C. The cell concentration for each region was estimated by mixing 10 µL of each sample with 1 µL of 100x SYBR Gold (Invitrogen, Cat no. S11494), pipetting 10 µL on a hemocytometer (Hausser Scientific, PA, USA) secured on a glass slide and counting cells at 5X magnification using an inverted LED fluorescence motorized microscope (Zeiss LSM 710 microscope, Carl Zeiss Microscopy, GmbH, Germany). Every sample was then brought to 300,000 cells/mL in 25 µL with PBS and used for the Comet assay.

### Quantification of DSBs DNA damage using the Comet assay

Neutral Comet assays were performed to quantify the amount of DSB in different young or old mouse brain regions (CBL and STR). In brief, dispersed cells were counted and adjusted to 300,000 cells/mL A 25 µL aliquot of each sample was mixed with 250 µl molten low-melting agarose (R&D Systems, Cat no. 4250-050-02). 40 µL of this mixture was pipetted onto a 3-well FLARE slide (R&D Systems, Cat no. 3950-075-02) and spread with the pipette tip. For the Neutral Comet, slides were immersed in cool neutral buffer (0.1 M Tris, 0.5 M Sodium Acetate, pH 9) for 30 min, before electrophoresis was performed at 21 V for 45 min at 4°C in 850 mL of neutral buffer. Slides were put in DNA Precipitation Solution (1 M Ammonium Acetate in 95% EtOH), then 70% EtOH for 30 min each. For Comet assays, slides were dried at 37°C for 10-15 min. 100 µL 1x SYBR Gold (Invitrogen, Cat no. S11494) were placed onto each well for 30 min in the dark, before being rinsed with distilled water. Slides were dried completely at 37°C, and fluorescence 5x5 tilescan (0.6 zoom) images of the comets were captured at 5X magnification using an inverted LED fluorescence motorized microscope (ZEN 2.1 SP3 FP3 (black) Zeiss LSM 710 microscope, Carl Zeiss Microscopy, GmbH, Germany). Comet images were analyzed using Trevigen comet analysis software (R&D Systems, MN, USA). We scored at least 200 cells per sample. We have calculated DSBs using tail length and tail moment with equivalent results (85). The average value of DNA in the comet tail (%) and tail moment were used as the parameters for estimating basal DSB levels.

### Preparation and culturing of primary glia from brain regions

Intact brains were collected from day 1-3 newborn pups (P1-3) of male C57BL/6J mice. Brain regions (hippocampus, cerebellum, cortex, and striatum) pooled from 4-8 pups were isolated in a solution of HBSS supplemented with 1mM L-glutamine, 1mM sodium pyruvate and 1x Non-Essential Amino Acids. These tissue suspensions were digested in 5 mL 0.025% Trypsin-EDTA (Gibco 25300056) for 20 min at 37°C with gentle rocking. Tissue pieces were pelleted (5 min, 300g, room temperature (RT)) and then gently triturated 20-30 times in pre-warmed astrocyte media (Neurobasal A base media (Thermo-Fisher #10888022), 10% FBS (JRS 43635), 2% B27 Supplement (Themo-Fisher #1504044), 25mM glucose, 2mM sodium pyruvate, 2mM GlutaMax, 1x non-essential amino acids (Quality Biologicals 116-078-721EA), 1x antibiotic/antimycotic (Gibco #15240062) using a 5 mL pipet, to dissociate into cells. Each cell suspension was tested for mycoplasma and negative cultures were passaged 3 times to expand cell number. Each passage was cultured for 6-10 days (at 37°C, 5% CO_2_) with media exchanges every 2-3 days. At the end of three passages (roughly 25 days in culture), some cultures developed traces of mycoplasma but the transfection results of these cultures did not differ significantly from the negative cultures. Astrocyte cell purity and homogeneity was established by immunofluorescent analysis using anti-GFAP antibody-Cy3 conjugate (Abcam ab49874).

### Transfections in primary glia cultures

Primary mouse glia was used live or thawed and cultured in prewarmed DMEM high glucose medium supplemented with 20% fetal bovine serum, GlutaMax, non-essential amino acids, plasmocin, normocure, and penicillin/streptomycin. For transfection experiments, 0.05 million cells were seeded per well in 12-well plates and allowed to attach overnight. All cells were passages once to achieve about 80% confluency. On the day of transfection, cell culture medium was changed to 1 ml OPTI-MEM, and the cells were transfected with 1 µg plasmid cocktails using lipofectamine 3000 (Thermofisher Scientific) according to the manufacturer’s protocol. The transfection medium was changed to cell culture medium at 4 hours post transfection. At 24 hours post transfection, the cells were washed with PBS, trypsinized, and filtered through a 40 µm strainer cap tubes for flow cytometry analyses.

### FM-HCR Reporter vectors

All reporter plasmids used for FM-HCR were pMax vector-based. They were engineered with site-specific DNA lesions as previously described^82–85^. For NHEJ reporter, pMax BFP with a ScaI recognition site (BFP-ScaI) was linearized with ScaI restriction and purified using phenol-chloroform extraction and ethanol precipitation method. For the NER reporter, pMax mPlum was irradiated with UV-C light at 800 J/m^2^ and purified using ethanol precipitation method. The mPlum UV reporter was validated by analytical digest with T4PDG enzyme. For BER, the pMax GFP THF reporter was used to report long-patch base excision repair activity, and, for MMR, the pMax GFP A:C MMR reporter was generated using the described protocol. Briefly, the base plasmid pMax- GFP C289T was nicked with Nt. BspQ1, digested with ExoIII to generate ssDNA, annealed with an oligo, that contains the mismatch. Subsequently the oligonucleotide was extended by an overnight incubation with DNA polymerase and ligase, cleaned up with T5Exo digestion, precipitated with PEG8000 solution, and purified using phenol-chloroform extraction and ethanol precipitation. The GFP AC MMR reporter was validated in TK6 MSH2^-/-^ and WT cells.

HR activity was reported using a new BFP HR assay, which was designed with reference to the reported pCX-NNX-GFP HR assay^85^. Briefly, three gene blocks were designed to generate a wild-type plasmid (2Nt-BFP), a donor plasmid (2Nt-D3BFP), and a plasmid (2Nt-D5BFP) with a ScaI restriction recognition site. Each gene block contains a NheI recognition site at the 5’ end a SacI recognition site were deleted at the 3’end. Adjacent to the NheI recognition site are two recognitions sites for Nt. BspQ1 (2Nt) that flank the recognition sites for MluI and ScaI, and all of which are upstream of the Kozak consensus sequence. For 2Nt-D5BFP, the last four base pairs of the Kozak sequence and the first 19 base pair from the 5’ end of BFP were deleted to generate a novel Pst1 recognition site and a truncated BFP sequence that renders the protein encoded by the plasmid non-fluorescent. For 2Nt-D3BFP, 99 base pairs were deleted from the 3’ end to generate a truncated BFP protein that renders the protein expressed by the plasmid non-fluorescent. The 2Nt-BFP, which serves as the positive control, is engineered to have a full length BFP sequence, and thus is fluorescent. To generate the three HR reporters, pMax vector and gene blocks were digested with NheI and SacI and purified using gel extraction (Monarch DNA gel extraction kit, NEB) for the linearized vector and column purification (Monarch PCR and DNA cleanup kit, NEB) for the gene blocks. The gene blocks were then cloned into the linearized pMax vectors and amplified in DH5α competent cells. Putative positive clones were selected by kanamycin resistance and validated by sequencing using pMax reverse primer. Individual reporters were amplified in 1L LB medium with kanamycin and extracted using PureLink endotoxin-free giga plasmid purification kit (Thermofisher Scientific). For generating the linearized 2Nt-D5BFP for HR assay, 2Nt-D5BFP was linearized with PstI and purified using phenol-chloroform extraction and ethanol precipitation method. Fluorescence of each vector was tested in cell lines using flow cytometry analyses and the recombinogenic events between the linearized 2Nt-D5BFP and 2Nt-D3BFP were validated in HR deficient cell lines.

### FM-HCR Reporter cocktails

Three reporter cocktails were prepared for FM-HCR analyses. Damaged reporter cocktail A is composed of 100ng of PstI-linearized 2Nt-D5BFP (HR), 100ng of pMax GFP THF (BER), and 100ng of pMax mPlum as transfection control. Damaged reporter cocktail C is composed of 100ng of GFP_AC reporter (MMR) and 100ng of pMax_ mPlum as transfection control. A final amount of 2Nt-D3BFP was added to each damaged plasmid cocktail so a total of 1400 ng plasmid DNA was used for transfection. A single undamaged reporter cocktail is composed of 1100ng of 2Nt- D3BFP, 100ng of each of BFP_ScaI, pMax_GFP, and pMax_mPlum.

### FM-HCR Analysis

Primary glia were transfected with sufficient efficiency (4-8%) to afford a robust analysis of DNA repair capacity by flow cytometry analyses. All transfected cells were analyzed by flow cytometry (Attune NXT Flow cytometer, Thermofisher Scientific) at 24-hours post transfection. Data analyses were performed as previously described^82–84^ to determine the DNA repair capacity of the primary glia. Briefly, for each transfection (n=5), the product of the fluorescent positive cell counts and mean fluorescence intensity of the positive cells in each gate was normalized to that in the gate for transfection control. Activity of each DNA repair pathway was reported as % reporter expression, which is the ratio between the normalized reporter expression in the damaged and the undamaged cocktail multiplied by 100. The activity level of each DNA repair pathway studied is positively correlated with the % reporter expression. FM-HCR assays report independent repair mechanisms and absolute reporter expression depends on factors specific to each assay precluding statistical comparisons between pathways. Thus, % reporter expression approximates the percentage of reporter plasmids that have been repaired by the pathway of interest, this approach provides a rough estimate of the relative activity of multiple pathways based on this metric. FM-HCR assays report independent repair mechanisms and absolute reporter expression and depend on factors specific to each assay, precluding statistical comparisons between pathways. The FM-HCR assays are highly validated, robust reporters of DNA repair activity of cells and reflected activity of the line. For example, using the same FM-HCR assays, a panel of human lymphoblastoid cell lines from apparently healthy individuals consistently yielded around 2% reporter expression for HR, as was observed in the glia, but had greater than 30% reporter expression for NHEJ^82–84^. At the same time, expression for the BER reporter was high (∼40%) in mouse brain glial cells and in human lymphoblastoid cell lines^82–84^.

### XJB-5-131 Treatment

XJB-5-131 (gift form P Wipf (University of Pittsburgh) was synthesized^61^ and treatment protocols were as previously described^8,62,85^. Lyophilized, powdered XJB-5-131 was reconstituted in DMSO at a concentration of 1 μg/μL. These samples were aliquoted and kept at -80°C. On the day of injection, the XJB-5-131 solution was mixed with 0.2 μm filtered and pre-warmed PBS (100°C) to reach a final concentration of 2 mg/kg mouse body weight in 200 μL solution. This was heated for 10 sec and the solution (200 μL) was injected, within 30min of preparation, by intraperitoneal injection (IP). Administration started at 60 weeks of age and continued three times per week for the length of the study. Vehicle treatments were identical except that XJB-5-131 was replaced by filtered PBS.

### Statistical analysis

Statistical analysis is reported as appropriate in each Figure. For all box and whisker plots, the boxes represent 50% of all data points with the line indicating the median value. The remaining 50% of datapoints are in the whiskers, distributed as the 25% maximal values above the box and 25% minimal values below the box. For CometChip assays, graphical representations were expressed as Mean ± SD and one-way ANOVA followed by Tukey’s multiple comparisons test was performed using GraphPad PRISM version 9.5.1 for Mac, GraphPad Software, San Diego, California USA. P<0.05 was considered as statistically significant using ANOVA models with post- *hoc* multiple comparison tests by GraphPad PRISM (GraphPad Software, LLC).

## Data Availability

There are no restrictions on the availability of these data. Source data are provided in Source Data file for plots reporting means/averages in bar charts and tables. Reagent used are presented in Supplementary Table 1. Antibody testing results are presented in Supplementary Table 1. Uncropped gel images are provided in Supplementary Source Data file.

## Acknowledgements

This work was supported by NIH (R01 NS060115 and R01 GM066359 to CTM and U01ES029520 to ZDN.

## Author Contributions

All authors participated in discussion of the data, data analysis, and contributed to the manuscript writing.

A.A.P. Prepared cells and tissue samples for all experiments, performed and single cell analysis throughout; MMR protein expression (Fig. 2, Supplementary Fig. 2, Supplementary Fig. 3 and Supplementary Fig. 4, Supplementary Fig. 5, Supplementary Fig. 6, Supplementary Fig. 7 and Supplementary Fig. 8, and DSB analysis in tissues (Fig. c,d) and in cells (Fig. 5c), IF of mouse tissues throughout the manuscript (Fig, 2, Fig. 3, Fig. 4 and Fig. 9); 8- oxoG assays (Fig.7b.c); XJB-5-131 *in vivo* analysis (Fig. 8c), and Cas10DA and Cas9 targeting and XJB-5-131 analysis in cells (Fig. 9).

A.C. Performed and analyzed the FM-HCR assay (Fig. 6)

J-H.Y. Performed and analyzed the western analysis of protein expression for MMMR recognition proteins (Fig.1 and Supplementary Fig. 1) and for other DNA repair pathways (Fig. 3).

G.B. Provided the brain sections from WT, *zQ175,* MSH3(-/-) and *zQ175/MSH3* or analysis of DSBs.

L.B. Performed and analyzed the neutral CometChip analysis in mouse tissue (Fig. 6),; Imaging and analysis of Fig 5E-G; performed IF, imaged & analyzed Fig 7 and Supp Fig 8.

Z.D.N. Overall direction of FM-HCR analysis and tools; participated in critical evaluation of the manuscript.

C.T.M. Overall experimental design and critical evaluation of the project; wrote the manuscript with contribution from all authors.

*Correspondence should be addressed to CTM or AAP

## SUPPLEMENTAL FIGURE LEGENDS

**Supplemental Fig. 1:**
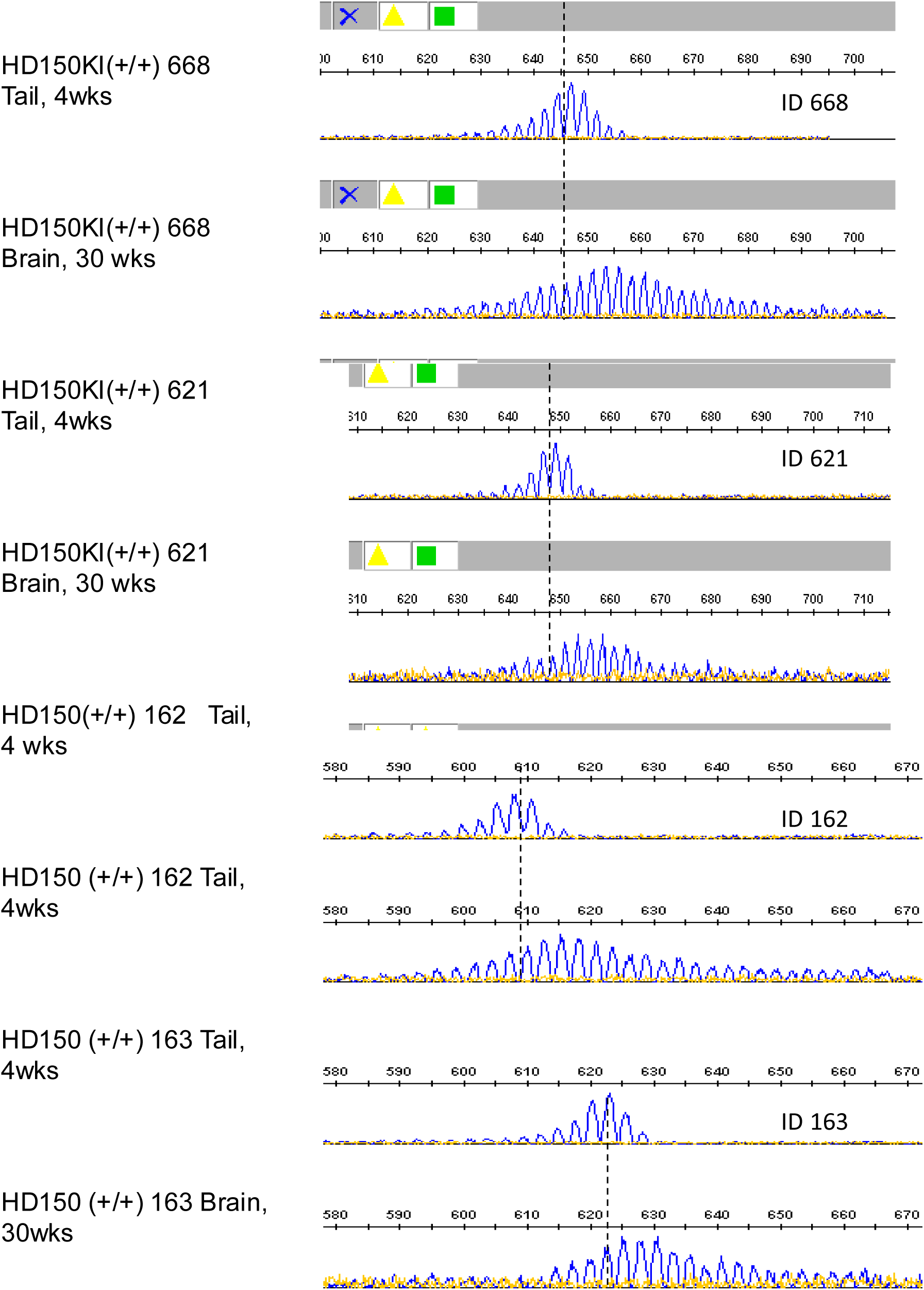
Somatic expansion occurs in the brains of *HdhQ(150/150*) mice and is longest in the STR. (A) The DNA was purified from the tails or tissue of animals was checked for quality by PCR. Good quality DNA was sent to TransnetYX CAG sizing by for high-resolution capillary electrophoreses. Representative examples of Genescan for somatic expansion in pairs showing the tail at 4wks and in the cortical brain tissue at 30 weeks in four *HdhQ(150/150*) littermates, IDs: 668, 621, 162, and 163. The dotted line indicates the midpoint of the tail a at 4wks, taken as the size of the inherited allele.

**Supplemental Fig. 2:**
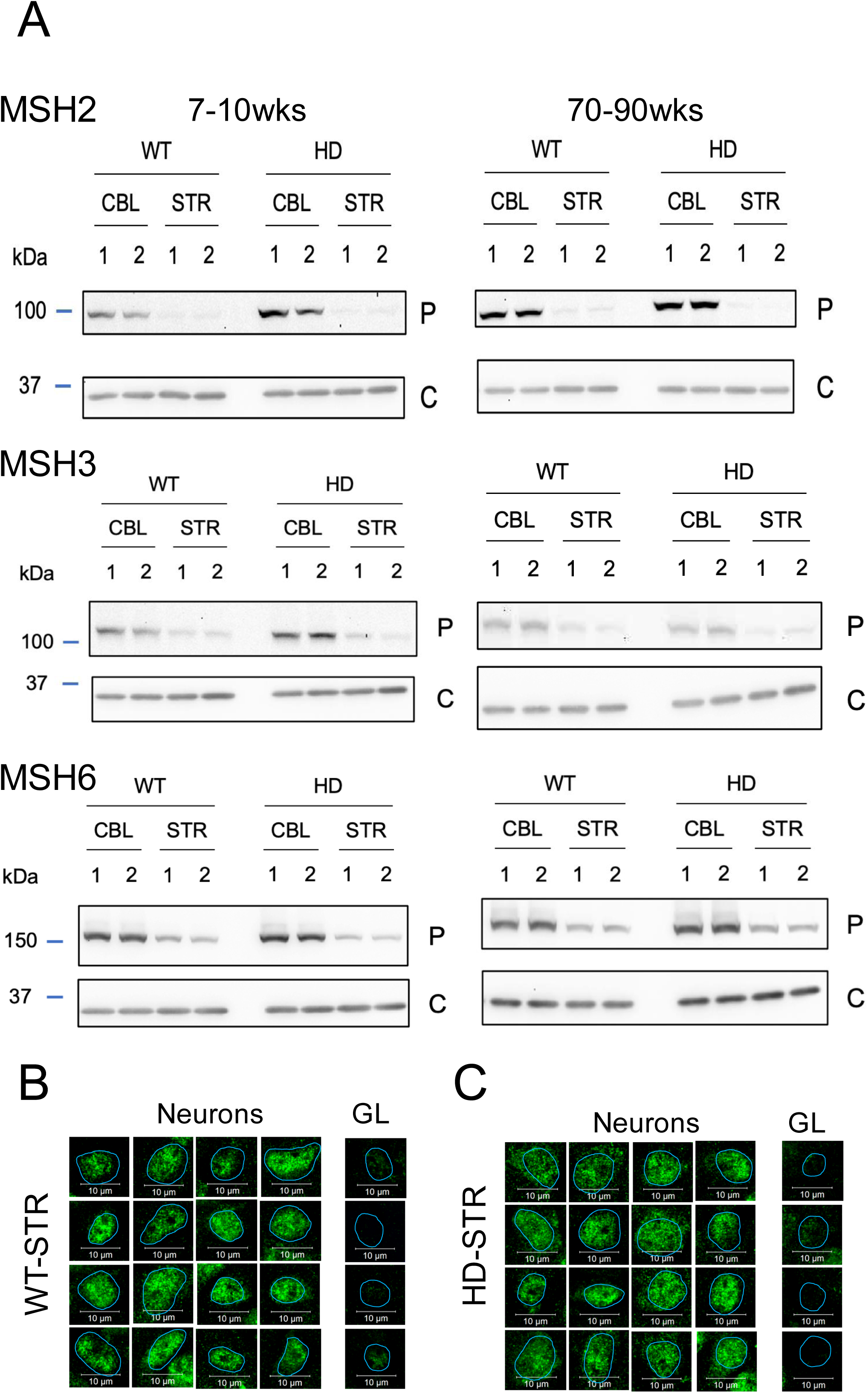
MSH3 and MSH6 expression is prominent in neurons and glia of WT and *HdhQ(150/150*) mice. (A) SDS-PAGE results for MSH2, MSH3 and MSH6 corresponding to the plots in Fig. 1. Details are described in the Fig. 1 legend. Protein extracts were resolved in six technical replicate SDS-PAGE gels; three gels for proteins measured in 10 wk animals (left) and three for proteins measured in 75 wk animals (right). Two siblings are shown side by side in each gel and are numbered as 1 and 2. Each blot was probed with indicated antibodies to a representative protein (P) or GAPDH (C). The protein expression data are normalized relative to the WT CBL. Error bars represent minimum (lower bar) and maximum (upper bar) values for the clonal samples at each age and were similar. The protein antibodies are listed in Supplementary Table 1. (B,C) Additional examples of magnified images of MSH3 as detected by MSH3 antibody IF staining (green) as shown in Fig, 2. Cell images were obtained from tissue sections from the STR of WT or HD animals of 70-90 wk. Neurons were NeuN (+) and GL were NeuN(-), as in Fig. 2. MSH3 expression is low in GL compared to neurons. The blue outline in the GL panels indicate the position of the nucleus as defined by DAPI staining. Scale bar is 10µm. The protein antibodies for MSH2, MSH3 and MSH6 are listed in Supplementary Table 2.

**Supplemental Fig. 3.**
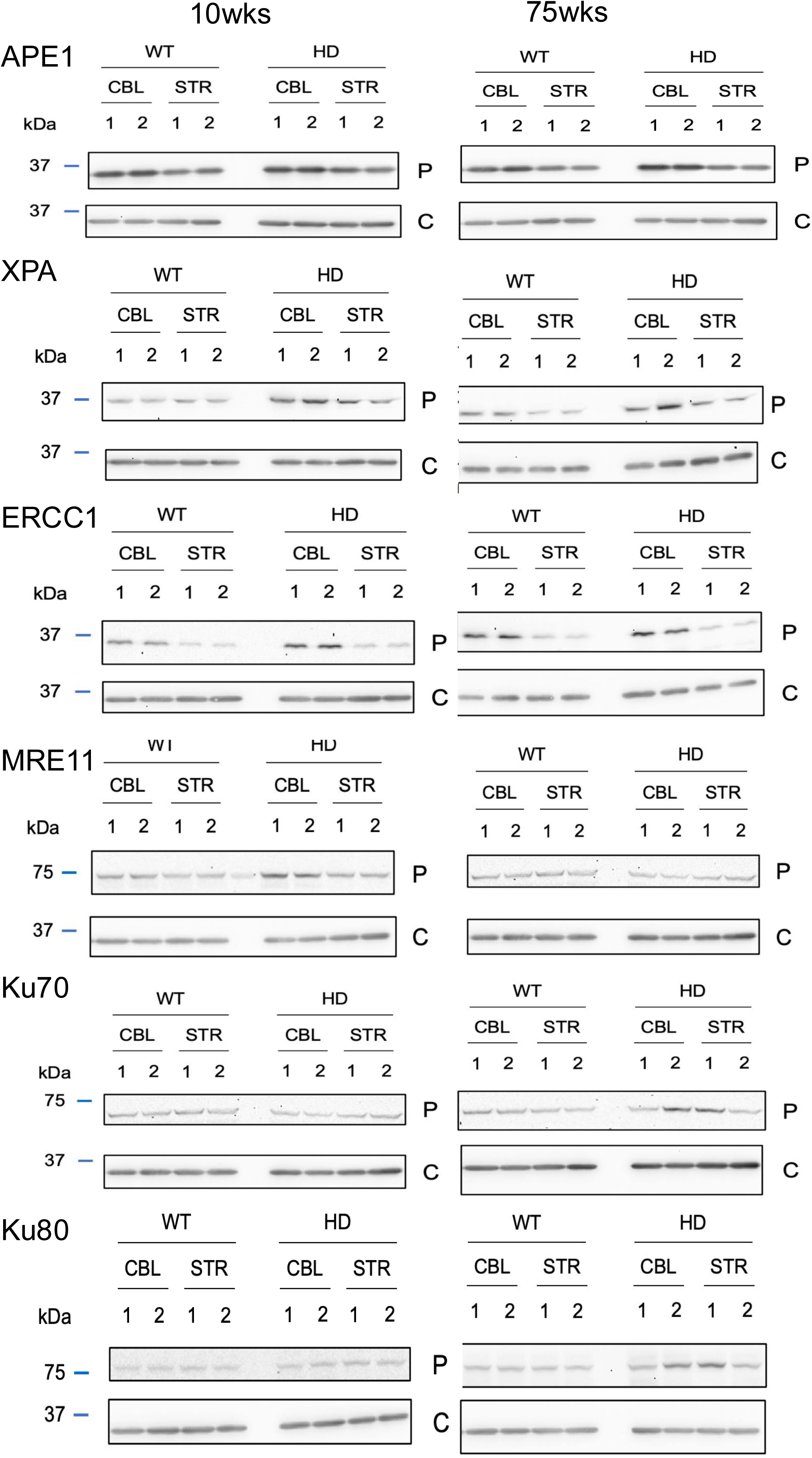
WT and *HdhQ(150/150*) mice express the machinery to carry out DNA repair. (A) SDS-PAGE results corresponding to the plots for DNA repair proteins in multiple pathways shown in Fig. 3. The representative pathway proteins measured in include Apurinic/apyrimidinic (AP) endonuclease (APE1); Xeroderma Pigmentosum Group A protein (XPA), Xeroderma Pigmentosum Group F protein (XPF), and Excision Repair Cross-complementation group 1 (ERCC1); Meiotic Recombination 11 Homolog 1 (MRE11); X-ray repair cross-complementing 6 (Ku70); X-ray repair cross-complementing 5 (Ku80). The proteins extracts were resolved in ten technical replicate SDS-PAGE gels; five gels for proteins measured in 10 wk animals (left) and five for proteins measured in 75 wk animals (right). Two siblings are shown side by side in each gel and are numbered as 1 and 2. Each blot was probed with indicated antibodies to a representative pathway protein (P) or GAPDH (C). The protein expression data are normalized relative to the WT CBL. Error bars represent minimum (lower bar) and maximum (upper bar) values for the clonal samples at each age and were similar. Source data and uncropped gels are provided in Supplementary Source File. The protein antibodies are listed in Supplementary Table 2.

**Supplemental Fig. 4:**
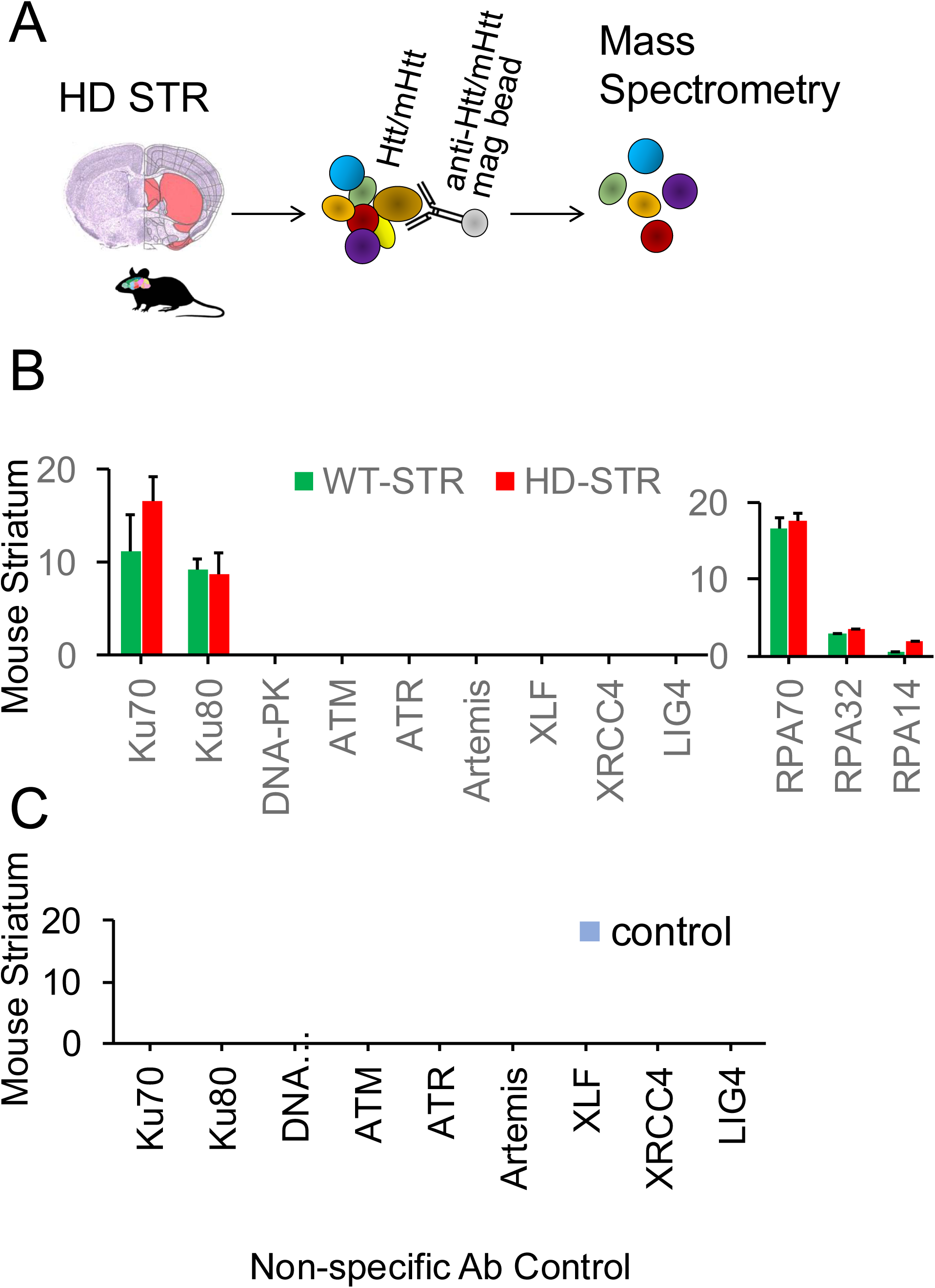
Htt and mhtt interacts with Ku70, Ku80 and RPA1-3 in the STR of *HdhQ(150/150*) mice. (A) Schematic diagram for immunoprecipitation and mass spectrometry (IP-MS) analyses of htt/Mhtt interactions with DNA repair proteins in the STR of WT or *HdhQ(150/150*) mice. The extracts from the STR of n=6 mice were immunoprecipitated using specific antibodies for htt and subjected to MS analysis to identify the binding partners. (B) Results for MS analysis of (A) for the pulldown products with the htt antibody for DNA repair proteins in the STR of WT (green) or *HdhQ(150/150*) (red) mice. Protein detected are Ku 70, Ku80 and RPA single strand binding proteins (right), which were like the MS results obtained in NIH3T293 cells (Fig. 4C,D). (C) No IP products were collected using a nonspecific antibody (control, blue). MS analysis was performed by the Mass spectrometry Facility at University of California, Davis (UC Davis).

**Supplemental Fig. 5.**
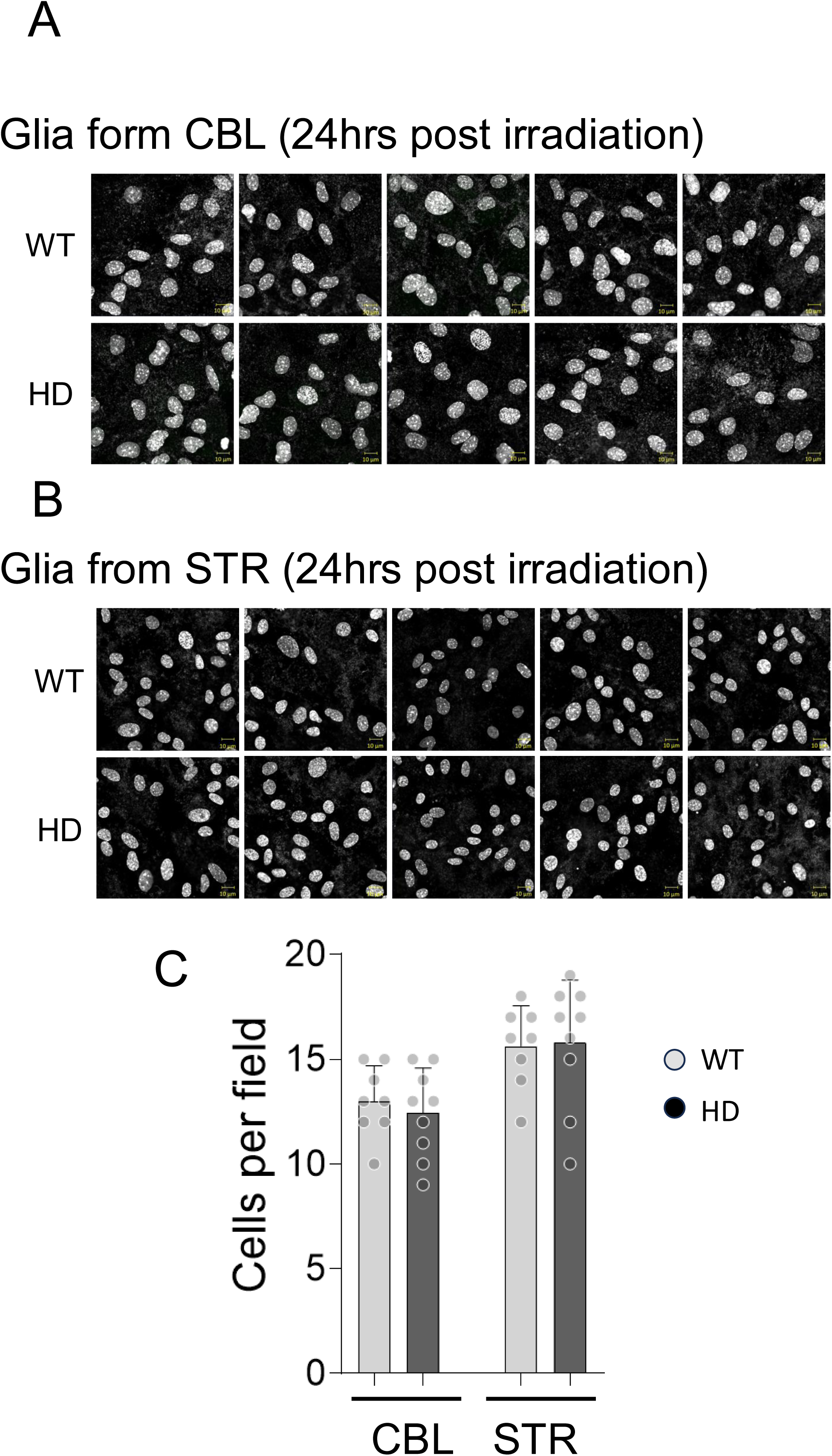
Radiation induced damage does not kill brain cells. (A,B) Purified glia were isolated from the CBL (A) or STR (B) of WT or *HdhQ(150/150*) animals and exposed to 2Gy radiation. Shown are images for n=5 random fields taken 24 hrs post irradiation. Scale bar is 10µm. The cells are intact and show good morphology. (C) Quantification of cell number in 10-20 random fields among n=3 platings. Cell counts were determined using a hemocytometer and the number was plotted for the STR and CBL from each genotype, WT (light gray) and HD (dark gray). There was no statistical difference among samples.

**Supplemental Fig. 6:**
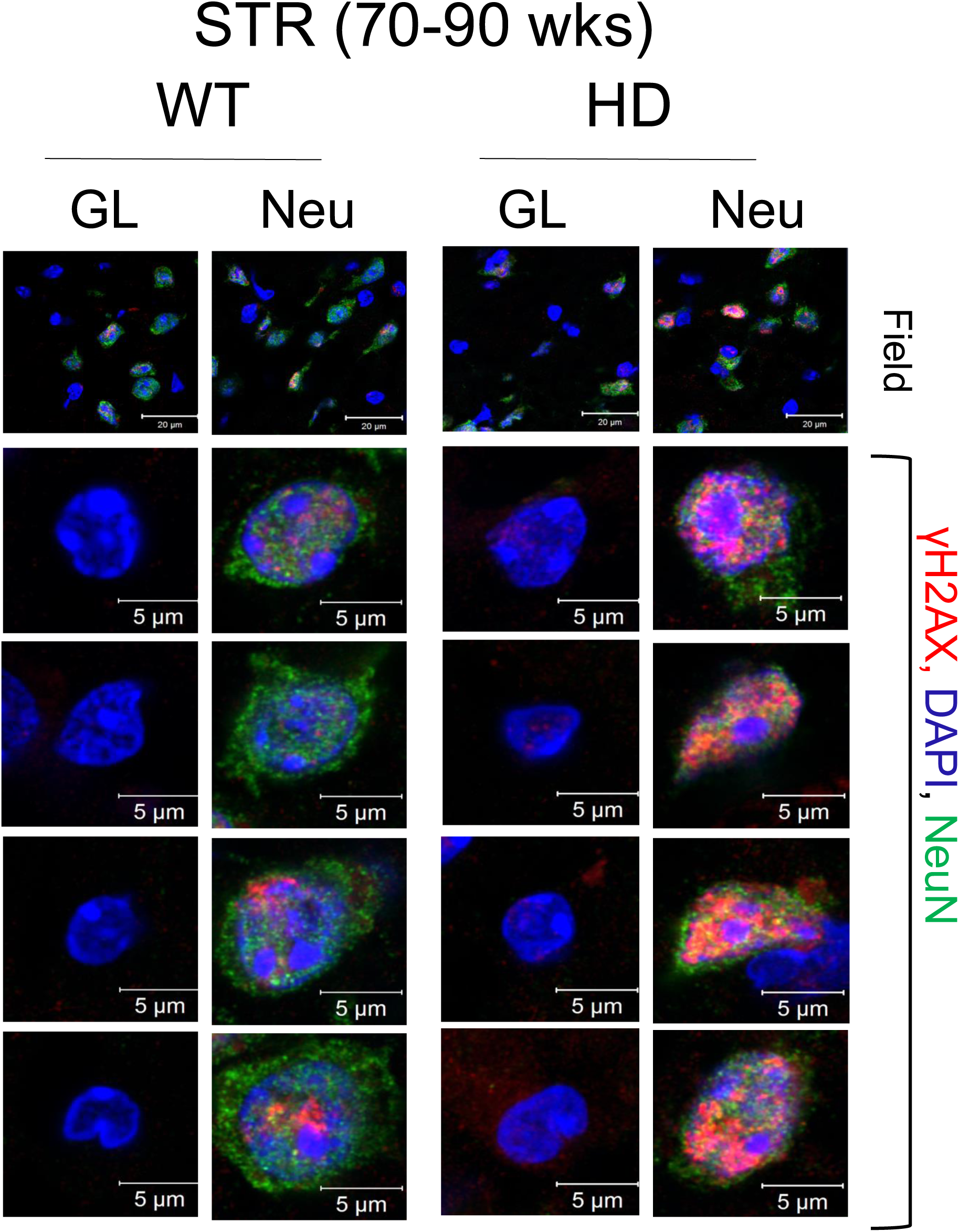
DSBs accumulate in neurons and Glia of WT and *HdhQ(150/150*) animals. Additional examples of images for DSBs measurements in tissue sections from the STR of 70- 90wk WT (left) or *HdhQ(150/150*)(right) animals, as described in Fig, 6 of main text. GL indicates glia and Neu indicates neurons. (Panel 1) An example of a random tissue field from the sections used for staining. The tissue sections were co-stained with nuclear DAPI (blue), neuronal NeuN (green) and DSBs marker γH2AX (red. Images shown are overlays of all three. (Panels 2-5) Individual magnified images of neurons and glia (GL) from the tissue sections. Scale bar is 5µm.

**Supplemental Fig. 7:**
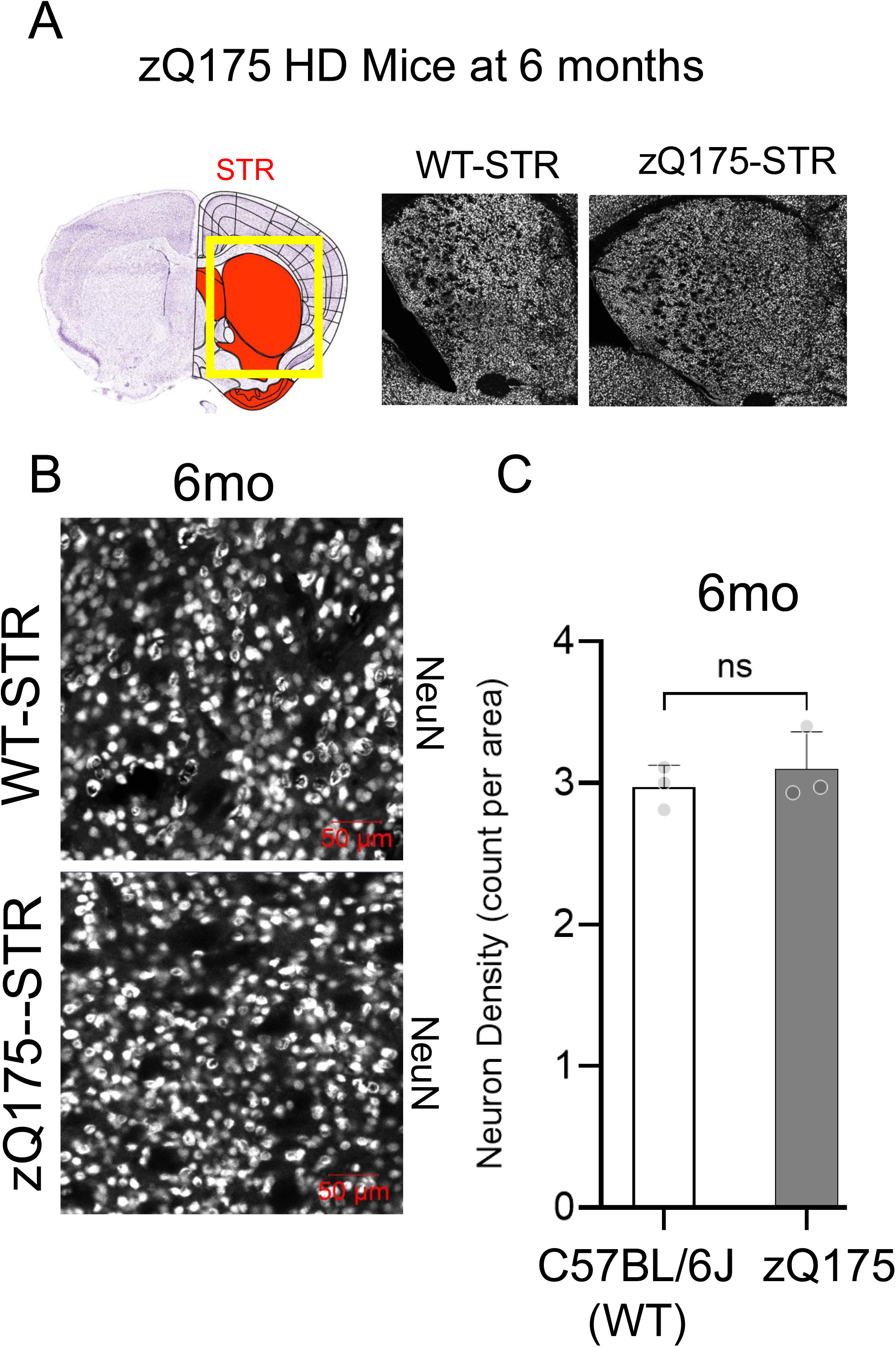
No loss of neurons in *zQ17 or zQ175/MSH3(-/-)* animals by 6 months. (A) (Left) Tissue section representation showing the position of the STR (red) and the tissue used for IF measurements (yellow box); (Right) images of tissue sections from the WT and *zQ175* mice at 6 months of age, as indicated. (B) Magnified images of tissue sections in (A) stained with an antibody to the neuronal marker, NeuN. Scale bar is 50µm. (C) Quantification of neuronal counts in tissue section from the STR of WT (left, white box) and zQ175 (right, gray box) animals at 6 months. The whole STR was imaged using ImageJ and the number of neurons per unit area was determined from the NeuN staining intensity using the software. The NeuN intensity was divided by the tile area to generate neuron density. Data taken from n=5 random fields from n=3 tissue slices from the STR of n=3 animals per genotype (WT vs HD). (ns) No statistical differences were observed in the neuronal counts.

**Supplemental Fig. 8:**
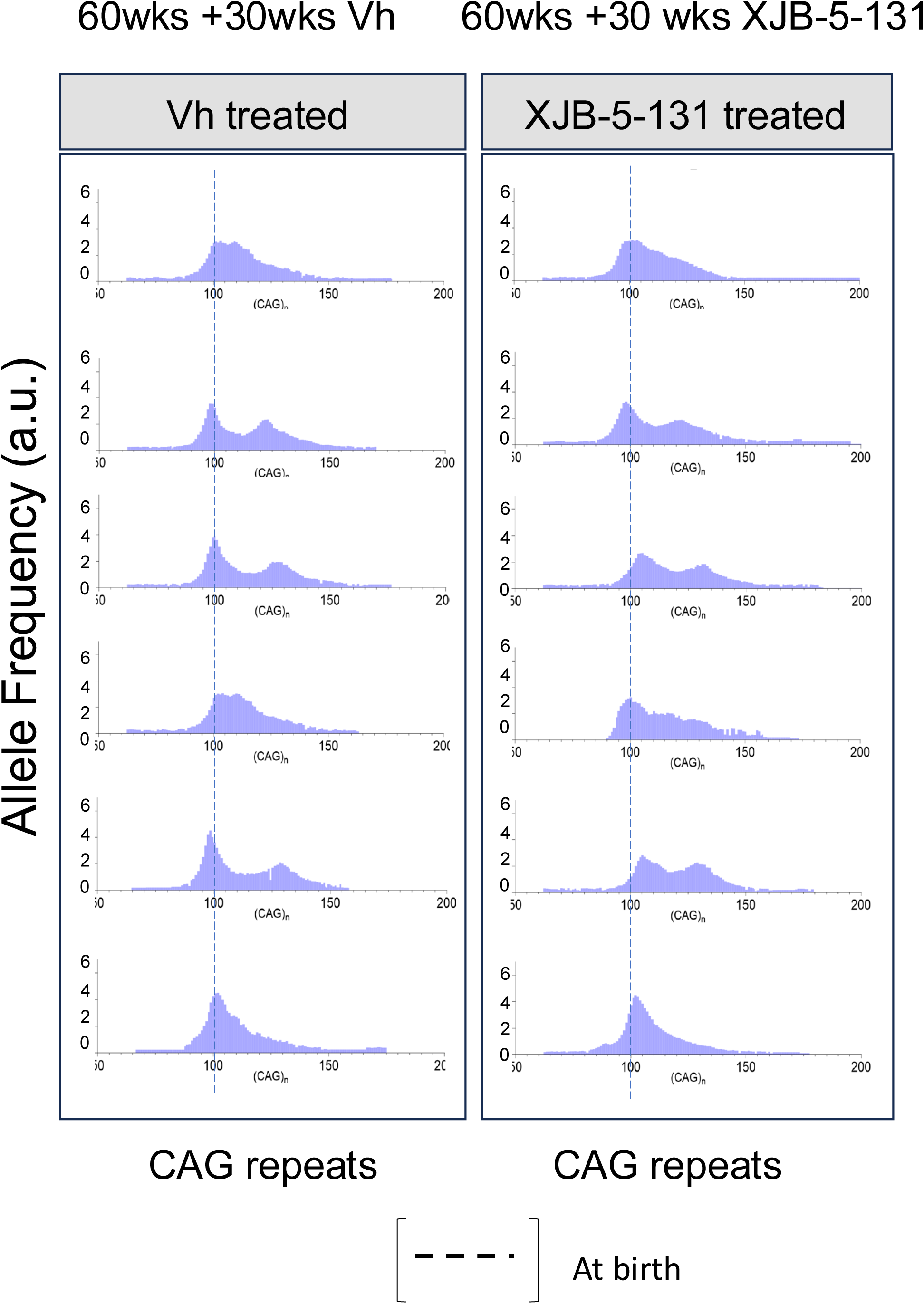
XJB-5-131 treatment of *HdhQ(150/150*) mice suppresses neuronal pathology with minimal effects on somatic expansion. The Genecan analysis of CAG repeat tracts of saline vehicle (Vh) or XJB-5-131 treatment at 90wks (60wks aging + 30 weeks treatment). Dotted line indicates CAG tract in tail at birth.

**Supplementary Table 1:** Key Resources.

| Reagent or Resource |  | Source | Catalogue ID |
| --- | --- | --- | --- |
| **Antibody** |  |  |  |
| Mouse anti-NeuN alexafluor-488 conjugate | 1:500 | EMD Millipore | MAB377X |
| Rabbit anti-NeuN alexafluor-647 conjugate | 1 :1,000 | Abcam | 190565 |
| Mouse anti-GFAP Cy3 conjugate | 1:500 | Abcam | ab49874 |
| Mouse anti-APE1 | 1:500 | Novus | 13B8E5C2 |
| Mouse anti-Ku80 |  | Santa Cruz | 515736 |
| Mouse anti-ERCC1 |  | Santa Cruz | 17809 |
| Rabbit anti-MSH2 |  | Abcam | ab92471 |
| Mouse anti-MSH3 |  | EMD Millipore | MABE324 |
| Rabbit anti-MSH6 |  | Abcam | Ab92471 |
| Rabbit anti-XPA |  | AbClonal | A1626 |
| Rabbit anti-MRE11 |  | Novus | NB100-142 |
| Mouse anti-yH2AX | 1:400 –<br>1:1,000 | ThermoFisher | MA1-2022 |
| Rabbit anti-53BP1 | 1:1,000 | Bethyl | A300-237A |
| Donkey anti-Mouse alexafluor-488 conjugate |  | Jackson Immunores. | 715-545-150 |
| Goat anti-Mouse alexafluor-568 conjugate |  | Invitrogen | A21124 |
| Donkey anti-Rabbit alexafluor-488 conjugate |  | Jackson Immunores. | 711-545-152 |
| Goat anti-Rabbit alexafluor-555 conjugate |  | Invitrogen | A32732 |
| Goat anti-Mouse alexafluor-488+ conjugate | 1:1,000 | Invitrogen | A48286 |
| Goat anti-Rabbit alexafluor-555+ conjugate | 1:1,000 | Invitrogen | A48283 |
| **Chemicals, peptides, and recombinant proteins** |  |  |  |
| Fc Receptor Block |  | Innovex | NB309 |
| Background Buster |  | Innovex | NB306 |
| Tissue-Tek O.C.T. Compound |  | Sakura | 4583 |
| TrueBlack |  | Biotium | 23007 |
| Monarch RNase A |  | New England Biolabs | T3018L |
| ImmuMount |  | Epredia | 9990402 |
| Nuclease P1 |  | New England Biolabs | M0660 |
| Quick Calf Alkaline Phosphatase |  | New England Biolabs | M0525 |
| CMV-cDNA26Q |  | constructed in house |  |
| CMV-cDNA51Q |  | constructed in house |  |
| T-PER Tissue Protein Extraction Reagent |  | Thermo Scientific | 78510 |
| Halt protease inhibitor cocktail |  | Thermo Scientific | 78420 |
| Pierce 660 Protein Assay reagent |  | Thermo Scientific | 22660 |
| NuPAGE Sample Reducing Agent |  | Invitrogen | NP0009 |
| Novex WedgeWell |  | Thermo Fisher | XP04205BOX |
| Amersham ECL Western Blotting Reagent |  | Sigma-Aldrich | GERPN2235 |
| CometAssay LMAgarose |  | R&D Systems | 4250-050-02 |
| CometAssay Lysis Solution |  | R&D Systems | 4250-050-01 |
| **Critical commercial assays** |  |  |  |
| DNA/RNA Oxidation Assay |  | Cayman chemicals | 589320 |
| DNeasy Blood and Tissue Kit |  | Qiagen | 69504 |
| **Experimental models:**<br>**Organisms/strains** |  |  |  |
| C57Bl/6J male mice |  | Jackson Labs | 000664 |
| NIH/3T3 fibroblasts |  | ATCC | CRL-1658 |
| **Software and algorithms** |  |  |  |
| ImageJ:Fiji |  | imagej.net/software/fiji/ | Version 2.15.0 |
| Prism Graph Pad |  | www.graphpad.com/features | version 9.5.1 |
| CometAssay Analysis Software |  | R&D Systems | 4260-000-CS |
| VersaDoc MP 4000 Imaging System |  | Biorad | Quantity One 1-D |
| Image Lab software |  | Biorad | V1 |
| ZEN Black |  | Zeiss | 2.1 SP3 FP3 |

**Supplementary Tabel 2.**
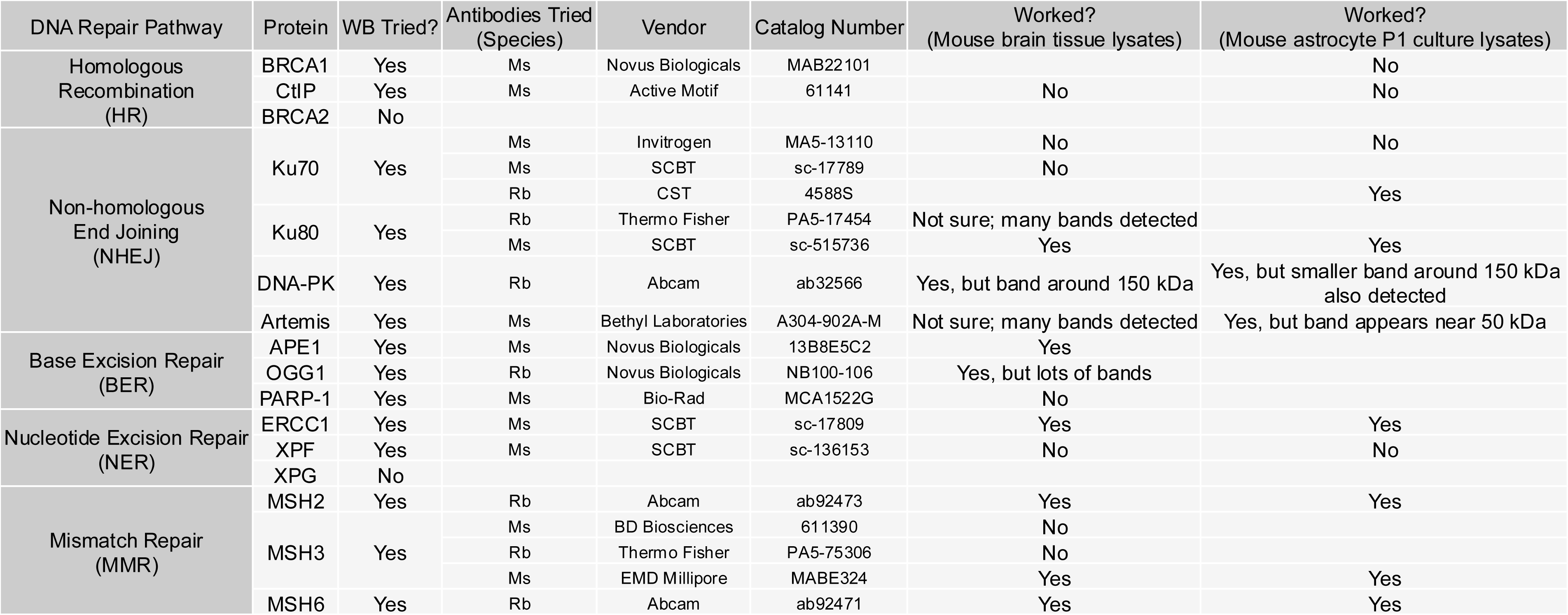
Antibody testing.

